# The olfactory basis of orchid pollination by mosquitoes

**DOI:** 10.1101/643510

**Authors:** Chloé Lahondère, Clément Vinauger, Ryo P. Okubo, Gabriella H. Wolff, Jeremy K. Chan, Omar S. Akbari, Jeffrey A. Riffell

**Affiliations:** Department of Biology, University of Washington, Seattle, WA, USA; Department of Biochemistry, Virginia Polytechnic Institute and State University, Blacksburg, VA, USA; Division of Biological Sciences, Section of Cell and Developmental Biology, University of California San Diego, CA, USA

**Keywords:** *Platanthera*, mosquitoes, *Aedes aegypti*, mosquito-plant interaction, nectar-feeding, pollination, olfaction

## Abstract

Mosquitoes are important vectors of disease and require sources of carbohydrates for reproduction and survival. Unlike host-related behaviors of mosquitoes, comparatively less is understood about the mechanisms involved in nectar-feeding decisions, or how this sensory information is processed in the mosquito brain. Here we show that *Aedes* spp. mosquitoes, including *Aedes aegypti*, are effective pollinators of the *Platanthera obtusata* orchid, and demonstrate this mutualism is mediated by the orchid’s scent and the balance of excitation and inhibition in the mosquito’s antennal lobe (AL). The *P. obtusata* orchid emits an attractive, nonanal-rich scent, whereas related *Platanthera* species – not visited by mosquitoes – emit scents dominated by lilac aldehyde. Calcium imaging experiments in the mosquito AL revealed that nonanal and lilac aldehyde each respectively activate the LC2 and AM2 glomerulus, and remarkably, the AM2 glomerulus is also sensitive to DEET, a mosquito repellent. Lateral inhibition between these two glomeruli reflects the level of attraction to the orchid scents: whereas the enriched nonanal scent of *P. obtusata* activates the LC2 and suppresses AM2, the high level of lilac aldehyde in the other orchid scents inverts this pattern of glomerular activity, and behavioral attraction is lost. These results demonstrate the ecological importance of mosquitoes beyond operating as disease vectors and open the door towards understanding the neural basis of mosquito nectar-seeking behaviors.

**Significance Statement:** Nectar-feeding by mosquitoes is important for survival and reproduction, and hence disease transmission. However, we know little about the sensory mechanisms that mediate mosquito attraction to sources of nectar, like those of flowers, or how this information is processed in the mosquito brain. Using a unique mutualism between *Aedes* mosquitoes and *Platanthera obtusata* orchids, we reveal that this mutualism is mediated by the orchid’s scent. Furthermore, lateral inhibition in the mosquito’s antennal (olfactory) lobe – via the neurotransmitter GABA – is critical for the representation of the scent. These results have implications toward understanding the olfactory basis of mosquito-nectar-seeking behaviors.

## Introduction

Mosquitoes are important vectors of disease, such as dengue, malaria or Zika, and are considered one of the deadliest animal on earth (*1*); for this reason, research has largely focused on mosquito-host interactions, and in particular, the mosquito’s sensory responses to those hosts (*2–6*). Nectar feeding is one such aspect of mosquito sensory biology that has received comparatively less attention, despite being an excellent system in which to probe the neural bases of behavior (*7*). For instance, nectar- and sugar-feeding is critically important for both male and female mosquitoes, serving to increase their lifespan, survival rate, and reproduction, and for males it is required for survival (*7, 8*).

Mosquitoes are attracted to, and feed from, a variety of plant nectar sources, including those from flowers (*9–13*). Although most examples of mosquito-plant interactions have shown that mosquitoes contribute little in reproductive services to the plant (*14*), there are examples of mosquitoes being potential pollinators (*10,11,15–18*). However, few studies have identified the floral cues that serve to attract and mediate these decisions by the mosquitoes, and how these behaviors influence pollination.

The association between the *Platanthera obtusata* orchid and *Aedes* mosquitoes is one of the few examples that shows mosquitoes as effective pollinators (*15–18*), and thus provides investigators a unique opportunity to identify the sensory mechanisms that help mosquitoes locate sources of nectar. The genus *Platanthera* has many different orchid species having diverse morphologies and specialized associations with certain pollinators (see *SI Appendix*, Table S1), with *P. obtusata* being an exemplar with its association with mosquitoes (*15–18*). Although mosquito visitation has been described in this species (*16*), the cues that attract mosquitoes to the flowers, and the importance of mosquito visitation for orchid pollination, are unknown.

In this article, we examine the neural and behavioral processes mediating mosquito floral preference. We present findings from (i) pollination studies in *P. obtusata* by *Aedes* mosquitoes, (ii) analyses of floral scent compounds that attract diverse mosquito species, and (iii) antennal and antennal lobe (AL) recordings showing how these floral scents and compounds are represented in the mosquito brain (Fig. S1). Using this integrative approach, we demonstrate that *Aedes* discrimination of *Platanthera* orchids is mediated by the balance of excitation and inhibition in the mosquito antennal lobe.

## Results

To understand the importance of various pollinators, including mosquitoes, on *P. obtusata*, we first conducted pollinator observation and exclusion experiments in northern Washington State where *Platanthera* orchids and mosquitoes are abundant. Using a combination of video recordings and focal observations by trained participants, more than 581 *P. obtusata* flowers were observed for a total of 47 h, with 57 floral feeding events by mosquitoes. During our observations, flowers were almost solely visited by various mosquito species (both sexes) that mainly belonged to the *Aedes* group (Fig. 1A,B; Table S2), with the only other visitor being a single geometrid moth. Mosquitoes quickly located these rather inconspicuous flowers, even on plants that were bagged and thus lacked a visual display. After landing on the flower, the mosquito’s probing of the nectar spur resulted in pollinia attachment to its eyes (Fig. 1A; Movies S1,2). Most of the pollinia-bearing mosquitoes had one or two pollinia, but we found up to four pollinia on a single female. To assess the impact of the mosquitoes’ visits on the orchid fruit set, we conducted a series of pollination experiments, such as bagging (thus preventing mosquito visitations) and cross- and self-pollinating the plants. We found significantly higher fruit-to-flower ratios and seed sets in unbagged plants compared with those in bagged or self-pollinated plants (Figs. 1C, S2; Mann-Whitney Test, p<0.001), and elevated fruit ratios in our cross-pollinated plants compared with bagged or self-pollinated plants (Fig. 1C). We then released field-caught mosquitoes into cages containing either a single plant or 2–3 plants (Fig. 1C,D). Once released into the cages, the mosquitoes fed from the *P. obtusata* flowers, and approximately 10% of the mosquitoes showed pollinia attachment (Fig. 1D). There was a trend for cages with two or more plants to have higher fruit-to-flower ratios than those with one plant (Mann-Whitney Test, p=0.07). Cages containing two or more plants had significantly higher fruit-to-flower ratios than bagged plants (Mann-Whitney Test, p<0.001), but were not statistically different from the unbagged plants (Mann-Whitney Test, p=0.84), further suggesting that cross-pollination is important in this orchid species.

**Figure 1.**
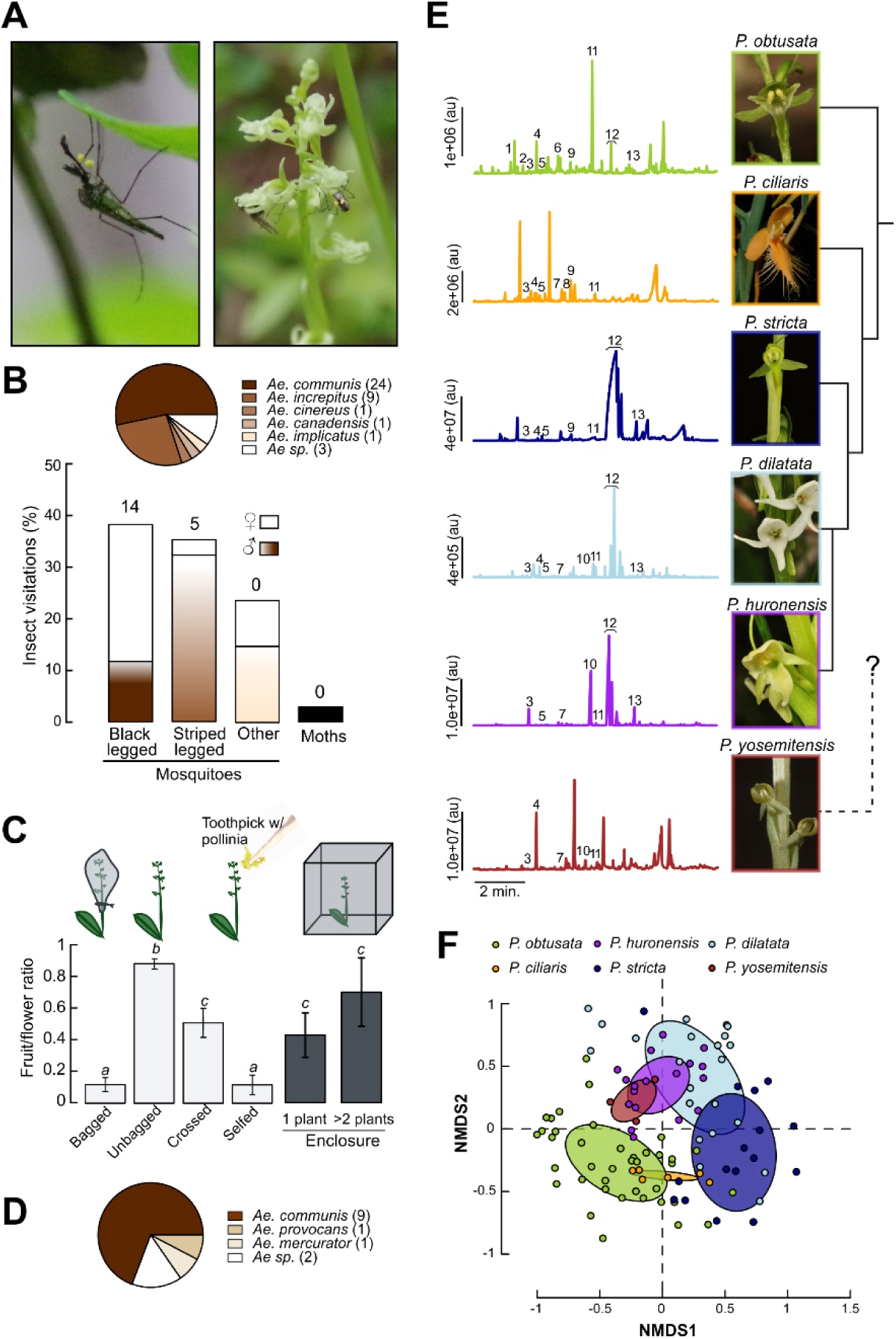
Association between the *P. obtusata* orchid and mosquito pollinators. (**A**) Picture (left image) of a black legged male mosquito bearing two pollinia on its head, and (right image) a male mosquito feeding on *P. obtusata* and a female with two pollinia attached to its head after having visited a flower. (**B**) Insect visitations (barplot; % insect visitation, calculated by the total number of insect visits to *P. obtusata*) and distribution of the mosquito species found in the field with pollinia (pie chart; numbers in legend denote the number of mosquitoes with pollinia). Both males (dark grey bars) and females (white bars) of different mosquito species visited the plants. Black-legged mosquitoes were pre-dominantly *Ae. communis*, and striped legged were *Ae. increpitus*. Numbers above the bars indicate the number of individuals observed with pollinia. (**C**) Fruit to flower ratio for bagged (using Organza bags around *P. obtusata* plants to prevent pollinator visitation), unbagged, self-crossed, out-crossed plants, and plants in the enclosure. Bagged and self-pollinated plants produced similar fruit-to-flower ratios (0.11 ± 0.04, 0.12 ± 0.06, respectively; Mann-Whitney Test: p = 0.99), but were significantly lower than the unbagged plants (0.89 ± 0.03; Mann-Whitney Test, p < 0.001). Although fruit weight did not differ between treatments (Student’s t-test, p = 0.082), bagged plants produced significantly less viable seeds per fruit per flower than unbagged plants (Fig. S1; Wilcoxon rank sum test, p < 0.05). Letters above bars show statistical differences between experimental conditions (Mann-Whitney Test: p<0.05). Bars are the mean ± SEM (n = 8-20 plants/treatment). (**D**) Pie chart of the species of mosquitoes which removed pollinia from the plants in the enclosures (numbers in legend denote the number of mosquitoes with pollinia). (**E**) Gas chromatography / mass spectrometry (GCMS) analyses of the floral volatiles emitted by *P. obtusata*, *P. ciliaris, P. stricta, P. dilatata, P. huronensis*, and *P. yosemitensis*. Pictures of the floral species, and their phylogenetic relationship, are shown on the right. *P. obtusata* flowers emitted a low emission rate scent that is dominated by aliphatic compounds (including octanal (#7), 1-octanol (#9), and nonanal (#11); 54% of the total emission), whereas the moth-visited species *P. dilatata, P. huronensis* and *P. stricta* emit strong scents dominated by terpenoid compounds (75%, 76% and 97% of the total emission for the three species, respectively), and the butterfly-visited *P. ciliaris* orchid is dominated by nonanal and limonene (24% and 12% of the total emission respectively) (SI Table 3). Numbers in the chromatograms correspond to: (1) α-pinene, (2) camphene, (3) benzaldehyde, (4) β-pinene, (5) β-myrcene, (6) octanal, (7) D-limonene, (8) eucalyptol, (9) 1-octanol, (10) (±)linalool, (11) nonanal, (12) lilac aldehydes (D and C isomers), and (13) lilac alcohol. Phylogeny to the right is from (*20*). (**F**) Non-metric multidimensional scaling (NMDS) plot (stress = 0.265) of the chemical composition of the scent of all the orchid species presented in B. Each dot represents a sample from a single individual plant collected in the field. The ellipses represent the standard deviation around the centroid of their respective cluster. Differences in scent composition and emission rate are significantly different between species (composition: ANOSIM, R=0.25, p=0.001; emission rate: Student t-tests, p<0.05).

**Figure 2.**
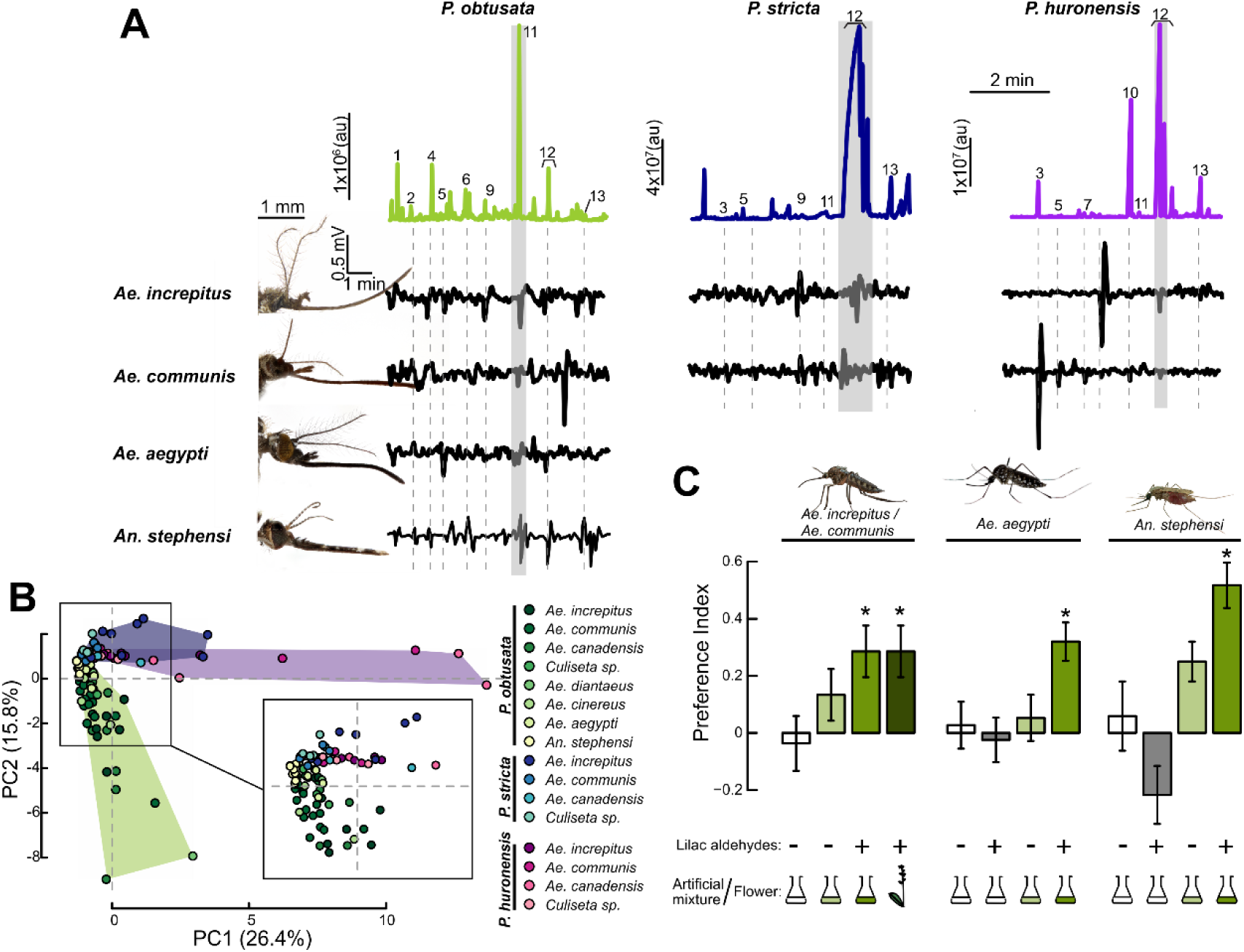
Identification of behaviorally effective orchid volatiles in mosquitoes. (**A**) Gas chromatogram traces for the *P. obtusata* (left), *P. stricta (*middle), and *P. huronensis* (right) headspaces, with electroantennogram responses to the GC peaks for four mosquito species (*Ae. communis, Ae. increpitus, Ae. aegypti*, and *An. stephensi*) immediately below. (1) α-pinene, (2) camphene, (3) benzaldehyde, (4) β-pinene, (5) β-myrcene, (6) octanal, (7) D-limonene, (8) eucalyptol, (9) 1-octanol, (10) linalool, (11) nonanal, (12) lilac aldehyde (C, D isomers), (13) lilac alcohol. (**B**) Principal Component Analysis (PCA) plot based on the antennal responses of individual mosquitoes from the different *Aedes* species to the peaks from the *P. obtusata, P. stricta* and *P. huronensi*s scents. Each dot corresponds to the responses of an individual mosquito; shaded areas and dots are color-coded according to mosquito species and flower scent (green, *P. obtusata*; blue, *P. stricta*; and purple, *P. huronensis*). Antennal responses to the three tested orchid scents were significantly different from one another (ANOSIM, R= 0.137, p < 0.01) (n=3-16 mosquitoes per species per floral extract). (**C**) Behavioral preferences by snow mosquitoes (*Ae. communis* and *Ae. increpitus*), *Ae. aegypti*, and *An. stephensi* mosquitoes to the *P. obtusata* scent and scent mixture, with and without the lilac aldehyde (at the concentration found in the *P. obtusata* headspace). A y-maze olfactometer was used for the behavioral experiments in which mosquitoes are released and have to fly upwind and choose between two arms carrying the tested compound / mixture or no odorant (control). A preference index (PI) was calculated based on these responses (see Supplementary Methods for details). The colored flask denotes the use of an artificial mixture (dark green is with lilac aldehyde; light green is without); empty flask denotes the negative (solvent) control. The plant motif is the positive control (orchid flowers), and the + and - symbols represent the presence or absence of the lilac aldehyde in the stimulus, respectively. Bars are the mean ± SEM (n = 27 - 75 mosquitoes/treatment); asterisks denote a significant difference between treatments and the mineral oil (no odor) control (binomial test: p<0.05).

### Platanthera orchids differ in their floral scents

*Platanthera obtusata* has a short (∼12 cm) inflorescence (Fig. 1A), and flowers emit a faint grassy- and musky-type of scent. The height and green coloration of the flowers make this plant difficult to pick out from neighboring vegetation, but over the course of our observations we noticed that mosquitoes readily oriented and flew to the flowers, exhibiting a zig-zagging flight typical of odor-conditioned optomotor anemotaxis (*6*). Moreover, even when the plants were bagged (thereby preventing the visual display of the flowers) mosquitoes would still land and attempt to probe the plants through the bag.

In the *Platanthera* genus, species differ in their floral advertisements, including their scent, and this is reflected in the different pollinators visiting each orchid species (Table S1). Often these species can co-occur in the same sedge, such as *P. obtusata*, *P. stricta, P. dilatata* and *P. huronensis*, although hybridization can be low (*19, 20*). Mosquitoes have sensitive olfactory systems that are used to locate important nutrient sources, including nectar (*2–5, 12*). Our observations on the strength of the association between *P. obtusata* and the mosquitoes, and how mosquitoes were able to locate the *P. obtusata* orchids, motivated us to examine the scent of closely related *Platanthera* species and identify the putative volatiles that mosquitoes might be using to detect and discriminate between the different orchid species.

The floral scents of the six orchid species were collected and subsequently characterized using gas chromatography with mass spectrometry (Fig. 1E). These analyses showed that species differed in both scent emissions and compositions (Fig. 1E,F; Table S3; composition: ANOSIM, R=0.25, p=0.001; emission rate: Student t-tests, p<0.05). Mosquito-pollinated *P. obtusata* flowers predominantly emitted nonanal and octanal, whereas the other orchid species, which are pollinated by other insect taxa (Table S1), emitted scents that were enriched in terpene compounds, such as lilac aldehyde (e.g., *P. dilatata, P. huronensis*, and *P. stricta*), or aromatic compounds, such as phenylacetaldehyde (e.g., *P. yosemitensis*).

### Divergent mosquitoes show similar antennal and behavioral responses to the *P. obtusata* orchid scent

To identify volatile compounds that mosquitoes might use to detect the plants, we performed gas chromatography coupled with electroantennographic detection (GC-EADs) using various species of mosquitoes that visit *P. obtusata* flowers in the field (Table S2). Several chemicals evoked antennal responses in the *Aedes* mosquitoes, including aliphatic (nonanal and octanal) and terpenoid compounds (e.g., lilac aldehydes, camphene and α- and β-pinene) (Figs. 2A, S3). For example. across the *Aedes-Ochlerotatus* group, nonanal elicited consistent responses and one of the strongest relative responses within a given mosquito species (Figs. 2A, S3). Interestingly, *Culiseta* mosquitoes, which also visited *P. obtusata* but did not have pollinia attachment, showed very little response to nonanal. Although mosquito species showed differences in their response magnitude to the chemicals (Figs. 2A, S3), the responses were relatively consistent which was reflected in their overlapping distribution in multivariate (Principal Components Analysis) space (ANOSIM, R = 0.076, P = 0.166)(Fig. 2B). This similarity in evoked responses by *Aedes* mosquitoes led us to examine whether these chemicals also evoked similar responses in other mosquitoes. We thus used two species of mosquitoes that are not native to the area, but are closely (*Ae. aegypti*) or distantly (*Anopheles stephensi*) related to the other *Aedes* species. The non-native mosquitoes (*Ae. aegypti* and *An. stephensi*) also responded to these volatiles and were not significantly different in their responses to the other *Aedes* species (ANOSIM, R = 0.087, p = 0.09)(Fig. 2B).

**Figure 3.**
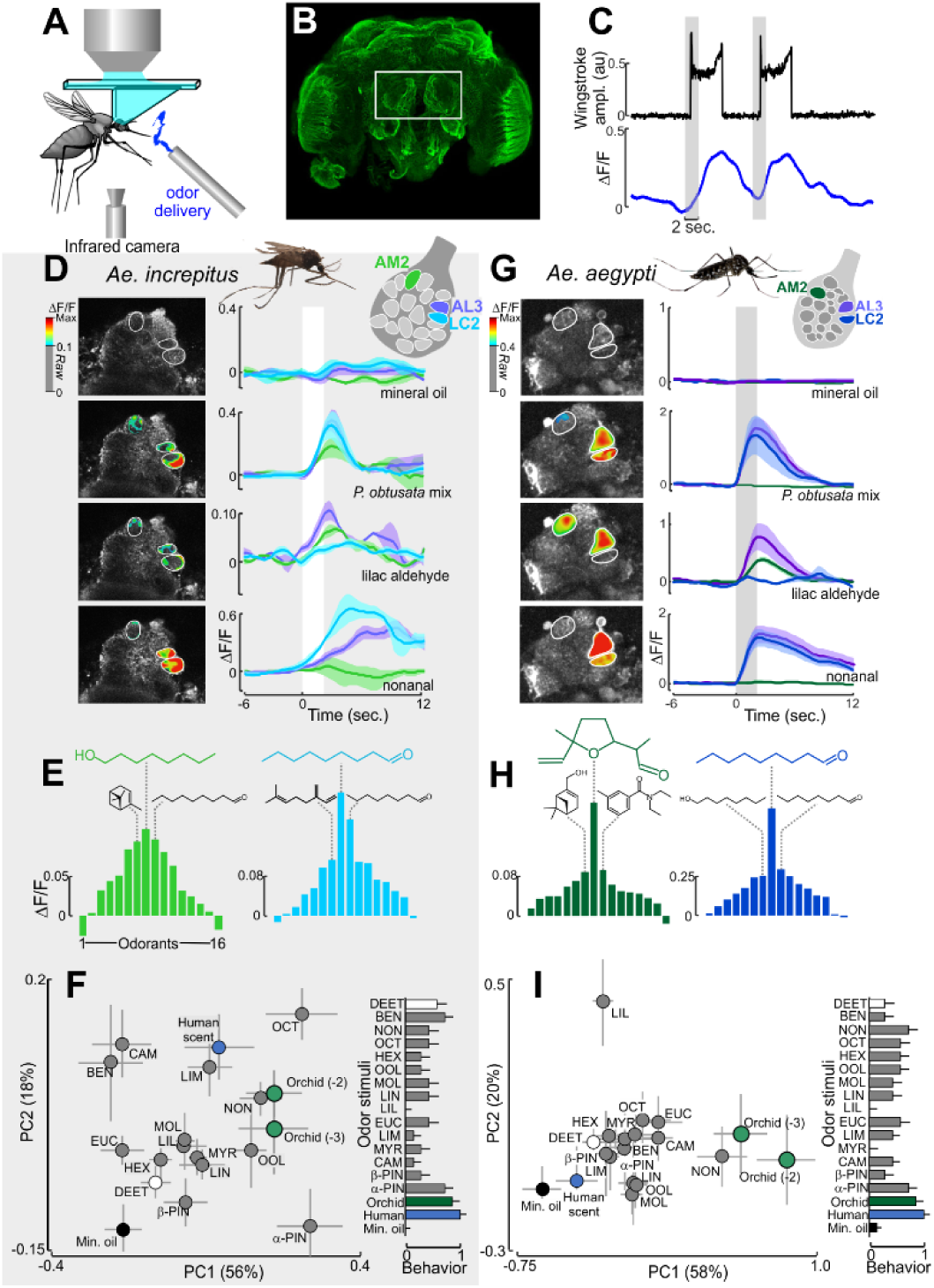
Mosquito antennal lobe responses to the *P. obtusata* scent. **(A)** Schematic of the two-photon setup used to record calcium dynamics in the mosquito AL. **(B)** *Ae. aegypti* brain (α-tubulin stain). The white rectangle surrounds the two ALs that are accessible for calcium imaging. Optical sectioning using the 2-photon microscope and subsequent immunohistochemical characterization allowed us to register glomeruli to an AL atlas as well as repeatably image from the same glomeruli. Although the AL between species differed in volume (0.0029±0.0001 and 0.0062±0.0004 mm^3^ for *Ae. aegypti* and *Ae. increpitus*, respectively), they consisted of similar numbers of glomeruli (18-22 glomeruli) in the ventral region of the AL, approximately 40 µm from the surface. **(C)** Representative time traces of behavioral (wing-stroke amplitude)(top, black) and AL LC2 glomerulus response (bottom, blue) to two *P. obtusata* odor stimulations (grey bars). **(D)** For *Ae. increpitus* mosquitoes with bath-application of Fluo4, schematic of AL glomeruli imaged at the 40 µm depth (top) and pseudo-color plot overlying the raw greyscale image (left) and mean ΔF/F time traces (right) for *Ae. increpitus* AL glomerular (AM2 [green], LC2 [blue] and AL3 [purple]) responses to mineral oil (no odor) control (top); *P. obtusata* mix (middle, top); lilac aldehyde (middle, bottom); and nonanal (bottom). White bars are the odor stimulations. Traces are the mean from 3-9 mosquitoes; shaded areas denote the SEM. Pseudo-color images were generated by subtracting the frame before stimulus onset from the frames during the stimulus window; only those glomerular regions of interest that were >0.1 ΔF/F are shown. **(E)** Tuning curves for the *Ae. increpitus* AM2 (green) and LC2 (blue) glomeruli based on a panel of sixteen odorants. AM2 is most responsive to octanol (green chemical structure), followed by α-pinene and nonanal (black chemical structures). LC2 is most responsive to nonanal (blue), followed by octanal and β-myrcene (black chemical structures). Bars are the mean (n=3-9). **(F)** (Left) Principal component (PC) plot from responses of 20 glomeruli to the odorants. PC1 and PC2 explain 56% and 18% of the variance, respectively. The orchid mixture at two concentrations (1:100 and 1:1000 dilution) and nonanal evoked stronger responses than the mineral oil (no odor) control (Kruskal-Wallis test: p<0.05) and were significantly different in the multivariate analysis (ANOSIM: p<0.05). Error bars represent SEM. (Right) Behavioral responses of the tethered mosquitoes to the odor stimuli. Responses were significantly different between the mineral oil control and the human and orchid scents (Kruskal-Wallis test: p<0.05), although they were not significantly correlated with the glomerular representations (Spearman rank correlation: *ρ*=0.35; p=0.16). **(G)** As in D, but for *PUb-GCaMP6s Ae. aegypti* mosquitoes and the AM2 (green), LC2 (blue) and AL3 (purple) AL glomeruli. Traces are the mean (n=7-14 mosquitoes); shaded area is the SEM. **(H)** As in E, but for the *Ae. aegypti* AM2 and LC2 glomeruli. AM2 is the most responsive to lilac aldehyde (green), followed by DEET and myrtenol (black chemical structures). LC2 is the most responsive to nonanal (blue), followed by octanal and octanol (black chemical structures). Bars are the mean (n=7-14 mosquitoes). **(I)** (Left) As in F, but for the *Ae. aegypti* mosquito and the 18 imaged glomerular responses to the panel of odorants. PC1 and PC2 explain 58% and 20% of the variance, respectively. (Right) Behavioral responses for the orchid and human scents were significantly different from control (p<0.05), although the correlation with the glomerular responses was not significant (Spearman rank correlation: *ρ*=0.46; p=0.07).

*P. obtusata* occurs in sympatry with *P. huronensis*, *P. dilatata* and *P. stricta*, but we did not observe *Aedes* mosquitoes visiting these orchids. To examine whether these differences in orchid visitation arise from differences in antennal responses, we performed GC-EADs using the scents of *P. stricta* and *P. huronensis*, which are predominantly pollinated by bees, moths, and butterflies (Table S1). Results showed that the mosquitoes (*Ae. increpitus, Ae. communis, Ae. canadensis*, and *Culiseta* sp.), which co-exist with these orchids in the same habitat, all responded to several compounds, including linalool, nonanal, benzaldehyde, β-myrcene and lilac aldehydes (Fig. 2). In particular, the high concentration of lilac aldehydes in the scent of *P. stricta*, and to a lesser extent in *P. huronensis*, elicited relatively strong responses in the antennae of *Ae. increpitus* and *Ae. communis*. Despite occurring in sympatry and overlapping in their scent composition, mosquito antennal responses to the three different orchid scents were significantly different from one another (Fig. 2B; ANOSIM, R= 0.137, p < 0.01), suggesting that the orchid species pollinated by other insects were activating distinct olfactory channels in the mosquitoes.

To evaluate if the *P. obtusata* orchid scent attracts mosquitoes, we tested the behavior of *Ae. increpitus* and *Ae. communis* mosquitoes (both important pollinators of *P. obtusata*) in response to the scent emitted by live *P. obtusata* flowers, as well as by an artificial mixture composed of the floral volatiles that elicited strong antennal responses in mosquitoes. Both the artificial mixture and the scent from the flowers significantly attracted these mosquitoes (Fig. 2C; binomial tests: p < 0.05). However, upon removal of lilac aldehyde (∼5.4 ng) from the mixture emissions, the attraction was reduced (binomial test: p = 0.292).

The similarity between mosquito species in their antennal responses to volatiles in the *P. obtusata* scent (Fig. 2) raised the question of whether closely related (*Ae. aegypti*) and more distantly related (*An. stephensi*) mosquitoes might also be attracted to the orchid scent. When tested in the olfactometer, both *Ae. aegypti* and *An. stephensi* mosquitoes exhibited significant attraction to the orchid scent with the lilac aldehydes (binomial tests: p<0.05). By contrast, and similar to responses by *Aedes* mosquitoes, once the lilac aldehydes were removed from the mixture this attraction was reduced to levels approaching the mineral oil (no odor) control (Fig. 2C). Nonetheless, the attraction by these other mosquito species may not indicate that pollinia also attaches to their eyes, or that they may serve as pollinators. To address this question, we released both male and female *Ae. aegypti* mosquitoes into cages with flowering *P. obtusata* plants. Once entering the cage, the mosquitoes immediately fed from the flowers, and pollinia attached to their eyes similar to the other *Aedes* species (Fig. S4).

### The *P. obtusata* orchid scent evokes strong responses in the mosquito antennal lobe

The differences in floral scents between the orchid species, and the behavioral responses by different mosquito species to the *P. obtusata* scent, raised the question of how this chemical information was represented in the mosquito’s primary olfactory center, the antennal lobe (AL). Therefore, we used bath application of a calcium indicator (Fluo4) in *Ae. increpitus* and our *PUb-GCaMP6s* line of *Ae. aegypti* mosquitoes (*21, 22*). Although both indicators of calcium (Fluo4 and *PUb-GCaMP6s*) do not allow explicit recording of specific cell types in the AL, but they do provide an ability to record and characterize the responses of individual glomeruli to odor stimuli. Mosquitoes were glued to holders that permitted two-photon imaging of calcium responses in the AL during tethered flight (*22, 23*) and tentative registration and naming of glomeruli (Fig. 3A,B). For both mosquito species, odor stimulation evoked distinct calcium dynamics in the glomerular regions of the AL that were time-locked to stimulus onset (Fig. 3C,D,G). The orchid mixture evoked flight responses and strong (>20% ΔF/F) multi-glomerular patterns of activity in both mosquito species, particularly in the anterior-medial glomeruli (the putative AM2, AM3, and V1 glomeruli) and the anterior-lateral glomeruli (AL3, and LC2) (Figs. 3D,G; S5, S6). In addition, certain odorants elicited overlapping patterns of glomerular activity similar to those elicited by the orchid scent (Fig. 3F,I), such as nonanal in the AL3 and LC2 glomeruli (Fig. 3D,G), with the LC2 glomerulus showing the strongest tuning to nonanal, octanal, and 1-octanol (Fig. 3E,H). Although the anterior-medial glomeruli showed broader tuning in *Ae. increpitus* than in *Ae. aegypti*, these glomeruli were sensitive to terpene compounds in both species and the AM2 glomerulus often exhibited inhibition when stimulated with nonanal (Figs. 3D,E,G, and H; S5, S6). Interestingly, the AM2 glomerulus showed the strongest tuning to lilac aldehyde, followed by DEET, a strong mosquito repellent (*24–27*)(Fig. S7), although these responses were suppressed when stimulated with the orchid mixture (Figs. 3G,H; S6). However, other odor stimuli, including human scent, evoked a dissimilar pattern of glomerular activity compared with the orchid mixture (Fig. 3F,I).

### Inhibition in the mosquito AL plays an important role in the processing of the orchid scents

Results from our calcium imaging and behavioral experiments suggested that certain volatile compounds, such as nonanal and lilac aldehyde, are particularly important for mosquito responses to *P. obtusata*. However, other *Platanthera* species, that are primarily pollinated by different insects, also emit these volatile compounds, but at different ratios (Fig. 4A), therefore raising the question of how mosquitoes respond to these scents. Behaviorally testing the scents of the moth- and bee-pollinated *Platanthera* orchids showed that these scents elicited behavioral responses that were not significantly different from the solvent control (binomial tests: p > 0.05), or elicited an aversive response when compared with the *P. obtusata* mixture (Fig. 4B; binomial tests: p < 0.05). To determine a correlation between mosquito behavior and AL response, we compared glomerular responses to the odors of the different orchid species. Stimulation with the *P. obtusata* mixture evoked strong glomerular responses in the AL, particularly in the AL3 and LC2 glomeruli, whereas stimulation with the other *Platanthera* scents (containing much higher lilac aldehyde: nonanal ratios) showed decreased responses in the LC2 glomeruli; however, the AM2 glomerulus (tuned to lilac aldehyde and DEET) showed much stronger responses (Figs. 4C,D; Kruskal-Wallis test with multiple comparisons: p < 0.05).

**Figure 4.**
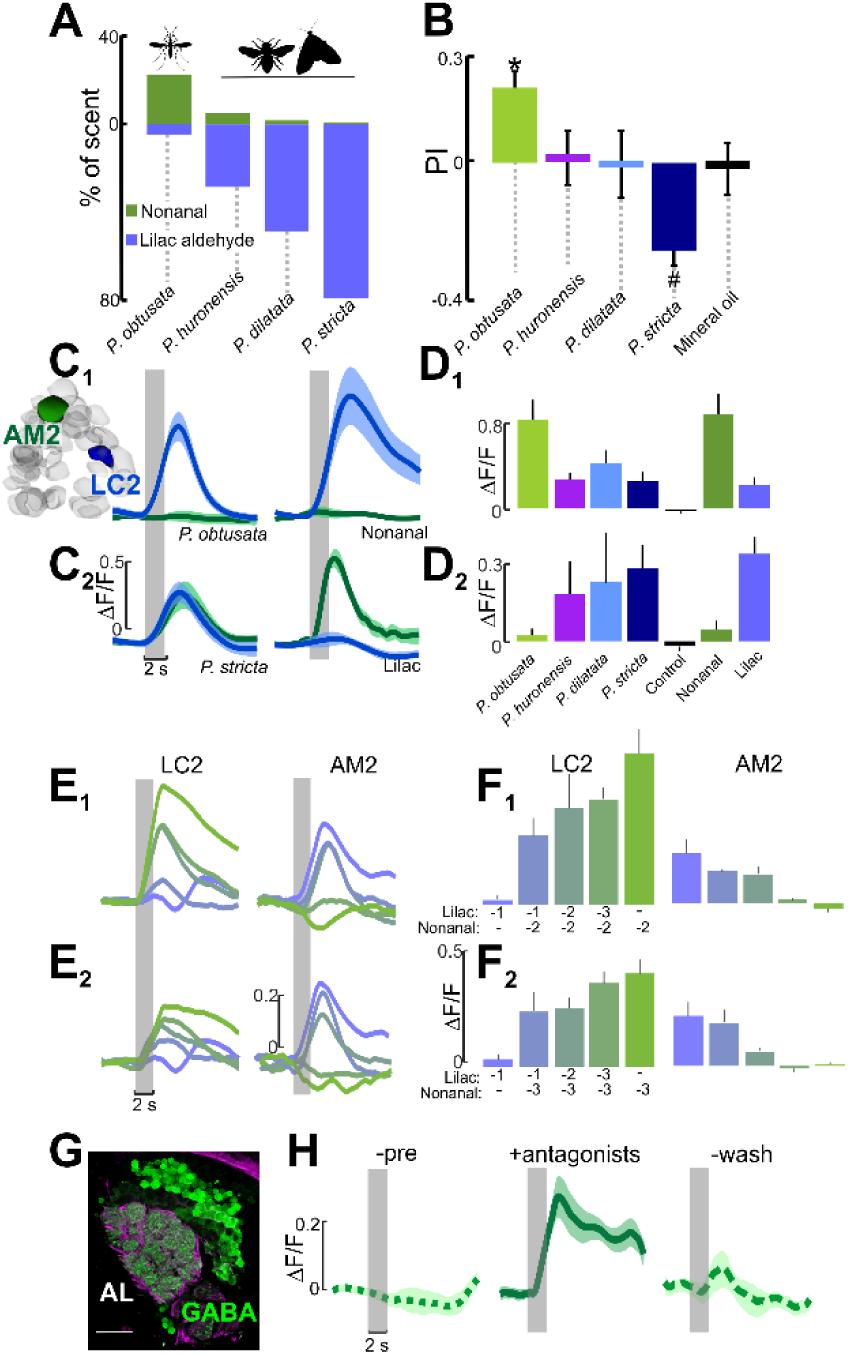
Glomeruli encoding the orchid scents are sensitive to odorant ratios. (**A**) Percentage of nonanal and lilac aldehyde concentrations in the different *Platanthera* orchid scents, which have 6- to 40-fold higher lilac aldehyde concentrations than *P. obtusata*. (**B**) Behavioral preferences by *Ae. aegypti* mosquitoes to scent mixtures containing lilac aldehydes at the concentrations quantified in the different *Plathanthera* species. Similar to Fig. 2C, mosquitoes were released in a y-olfactometer and had to choose between two arms carrying the scent mixture or no odorant (control). Asterisk denotes a significant difference from the mineral oil control (binomial test: p<0.05); number symbol denotes a significant difference from the *P. obtusata* scent (binomial test:p<0.05). (**C**) (C_1_) Mean ΔF/F time traces for LC2 (blue) and AM2 (green) glomeruli to *P. obtusata* (left) and nonanal (right). (C_2_) Same as in C_1_, except to the *P. stricta* scent (left) and lilac aldehyde (right). The *P. obtusata* and *P. stricta* mixtures contain the same concentration of nonanal and other constituents but differ in their lilac aldehyde concentrations (see panel A). Traces are the mean (n=6-10 mosquitoes); shaded areas denote ±SEM. (**D**) (D_1_) Responses of the LC2 glomerulus to the different *Platanthera* orchid mixtures, and the single odorants nonanal and lilac aldehyde. The increasing concentration of lilac aldehyde in the other orchid mixtures caused a significant suppression of LC2 response to the nonanal in the scents (Kruskal-Wallis test: p<0.05), even though nonanal was at the same concentration as in the *P. obtusata* mixture. (D_2_) Responses of the AM2 glomerulus to the different *Platanthera* orchid scents and nonanal and lilac aldehyde constituents. The increasing concentration of lilac aldehyde in the other orchid scents caused a significant increase in AM2 responses compared with responses to *P. obtusata* (Kruskal-Wallis test: p<0.05). Bars are the mean ± SEM. (**E**) ΔF/F time traces for the LC2 (left) and AM2 (right) glomeruli. The preparation was simultaneously stimulated using separate vials of lilac aldehyde and nonanal at different concentrations to create 10 different mixture ratios. (E_1_) Each trace is a different ratio of lilac aldehyde to nonanal, ranging from green (10^-2^ nonanal: 0 lilac aldehyde) to purple (0 nonanal: 10^-1^ lilac aldehyde); 10^-3^ to 10^-1^ lilac aldehyde, and 10^-2^ nonanal concentrations were tested. (E_2_) As in E_1_, except tested concentrations were 10^-3^ to 10^-1^ for lilac aldehyde, and 10^-3^ for nonanal. (**F**) (F_1_) Mean ΔF/F during 2 sec. of odor presentation for the LC2 glomerulus (left) and the AM2 glomerulus (right). Bars are color-coded according to the ratio of lilac aldehyde to nonanal traces in E_1_. (F_2_) As in F_1_, except the concentrations of lilac aldehyde and nonanal in the ratio mixtures correspond to those in E_2_. Bars are the mean (n=6) ± SEM. (**G**) Antibody labeling against GABA (green) in the right *Ae. aegypti* AL; background label (alpha-tubulin) is purple. Scale bar is 20 µm. (**H**) Mean ΔF/F time traces for the AM2 glomerulus. GABA receptor antagonists block the suppressive effect of nonanal to AM2’s response to the lilac aldehyde in the *P. obtusata* mixture, causing a significantly higher response than the pre-application and wash periods (Kruskal-Wallis test: p<0.05). Traces are the mean (n = 4 mosquitoes) ± SEM.

To better understand how the ratio of lilac aldehyde and nonanal altered the activation of the LC2 and AM2 glomeruli, we tested mixtures of lilac aldehyde and nonanal at different concentration ratios and found that lilac aldehyde suppressed the response of LC2 to nonanal, suggesting lateral inhibition between these two glomeruli. Higher lilac aldehyde concentrations increased LC2 suppression, but reciprocally increased AM2 activation (Fig. 4E,F). By contrast, nonanal caused suppression of AM2 responses to lilac aldehyde, with higher nonanal concentrations causing increased AM2 suppression, while increasing the activation of LC2 (Fig. 4E,F). To determine whether this suppression of glomerular activity is mediated by γ-aminobutyric acid (GABA), an important inhibitory neurotransmitter in insect olfactory systems (*28–30*), we used antisera against GABA in the *Ae. aegypti* brain and found widespread labelling in AL glomeruli, including AM2 and LC2 (Fig. 4G). Next, we pharmacologically manipulated the inhibition by focally applying GABA-receptor antagonists (1 µM CGP54626; 10 µM picrotoxin) on to the AL during our experiments. During application of the vehicle (saline) control, LC2 and AM2 responses to the *P. obtusata* scent were similar to those described above (Fig. 4E,F,H; S8), whereas during antagonist application, the effect of nonanal was blocked and the small amount of lilac aldehyde in the scent was sufficient to evoke a strong response in AM2 (Fig. 4H). The antagonists blocked the symmetrical inhibition by nonanal and lilac aldehyde in the *P. stricta* scent, causing increased response in both glomeruli, with the LC2 response levels similar to those evoked by *P. obtusata* (Fig. S8). Taken together, these results support the hypothesis that the ratios of volatile compounds in the orchid scents, and the resulting balance of excitation and inhibition in the mosquito AL, play an important role in mediating mosquito attraction to *P. obtusata* and possibly, reproductive isolation between orchid species.

## Discussion

In this study, we use a unique mutualism between *P. obtusata* orchids and *Aedes* mosquitoes to show the importance of mosquito pollination for this orchid and the role of scent in mediating this association. Olfactory cues play important roles in a variety of biological processes for mosquitoes, including locating suitable hosts (*5*), oviposition sites (*31*), and nectar sources (*32*). For *Aedes* mosquitoes to efficiently locate sources of nutrients, they must distinguish between complex floral scents in a dynamic chemical environment (*7*). In the case of sympatric *Platanthera* orchids – which share the same scent constituents but differ in their ratios of nonanal and lilac aldehydes –, their scents evoke distinct patterns of activation in AL glomeruli. How is this occurring? Our results suggest that GABA-mediated lateral inhibition from the LC2 glomerulus that encodes nonanal (found in higher abundance in *P. obtusata*) suppresses responses of glomeruli encoding lilac aldehydes (abundant in the scent of the other *Platanthera* species) which allows mosquitoes to distinguish between orchids.

There are only a handful of mosquito-pollinated flowers, but some of these species have been shown to emit similar volatile profiles as *P. obtusata* (*8, 9, 32–34*). Our results showed that certain terpene volatiles, like lilac aldehyde, were important in the discrimination of the *P. obtusata* scent, and at low concentrations this volatile was important for attracting diverse mosquito species. In other mosquitoes, oxygenated terpene compounds that are derivatives of linalool, like lilac aldehyde and linalool oxide, were shown to elicit attraction to nectar sources (*13, 35, 36*). The qualitative similarities in the scent profiles of attractive nectar sources, and the attractiveness of the *P. obtusata* scent across mosquito species, raises the question of whether flower scents may be activating conserved olfactory channels, such as homologous odorant receptors (*35*). This will hopefully motivate research to identify the odorant receptors that are responsive to floral compounds, and their projections to the AL, such as the LC2 and AM2 glomeruli (*35*).

Our results also demonstrate the importance of mixtures and the processing of odorant ratios in *Aedes*. Interestingly, some of the volatile compounds emitted from blood hosts also occur in the *P. obtusata* scent, including nonanal (*37, 38*). However, in both *Ae. increpitus* and *Ae. aegypti* mosquitoes, the AL representations of host and orchid scents were different, suggesting that these odors may be processed via distinct olfactory channels. Despite the different glomerular ensemble responses, the complex nectar and host odors may share some of the same coding processes by AL circuits, including lateral inhibition of glomeruli. Similar to floral scents, human odors are complex mixtures that can differ between individuals in their constituent ratios, which may explain why mosquitoes often show behavioral preferences for certain individuals over others (*5, 39*). These dissimilarities have important epidemiological implications for disease transmission (*5, 40, 41*), and could be related to the subtle differences in the ratios of key compounds in an individual’s scent (*39*). Future work may explore if mosquito AL circuits process other complex odors, like those of human scent or other nectar sources, in a manner similar to that of the orchid scents, and whether the identified odorants and corresponding glomerular channels and modulatory systems can be leveraged in control interventions.

## Materials and Methods

Procedures for floral VOCs collection and analysis, mosquito rearing, the preparation used for GC-EAD experiments, behavior experiments and associated stimuli, olfactory stimuli and pharmacological reagents used in calcium imaging experiments, and immunohistochemistry are described in *SI Appendix*, **Supplementary Methods**.

### Orchid-pollinator observations and pollination experiments

#### Flower observations

Pollinator activity was monitored in the Okanogan-Wenatchee National Forest (47.847° N, 120.707° W; WA, USA) from late June to early July in 2016 and 2017 when the flowers of *P. obtusata* were in full bloom. Multiple direct and video observations of varying lengths from 30 minutes to 2.5 h were made for a total of 46.7 hours (15 hours of direct and 31.7 hours of video recordings). The observations were conducted from 10am to 8pm when mosquitoes were found to visit the flowers. Observations were recorded by visually inspecting each plant, with the trained observer approximately 1 m away from the plant – this distance did not influence the feeding and mosquito-flower visitation since no mosquito took off from the plant in the field and instead remained busy feeding from flower after flower. To further prevent the potential for observer interference, video observations were made using GoPro® Hero4 Silver (San Mateo, CA USA) fitted with a 128gb Lexar® High-Performance 633x microSD card. Videos were set at 720p resolution, 30 frames per second, and “Narrow” field of view. These settings were optimized for the memory capacity, battery life, and best resolution by the camera. Both observation methods, direct and video, provided similar visitation rates. The visitation time, insect identity, leg color and sex (for mosquitoes), were recorded from both direct and video observations. The number of feeding (defined by the probing into the flower using the proboscis) and visits (non-feeding or resting) were quantified per hour per flower for each pollinator type. Over the course of the experiments and observations, temperatures ranged from 9.6° to 32.3°C, with a relative humidity range of 13.4% to 100% (iButtons; Maxim Integrated™, San Jose, CA, USA, #DS1923). These experiments, therefore, captured both sunny and rainy weather conditions that were common in this area at this time of the year.

#### Pollinator addition experiments

To evaluate the contribution of mosquitoes to the pollination of *P. obtusata* orchids, we performed pollinator addition experiments during June through July in 2016. Mosquitoes were collected from the Okanogan-Wenatchee National Forest using CDC Wilton traps baited with carbon dioxide (John W. Hock Company, Gainesville, FL, USA). Carbon dioxide traps provide a standardized method to sample the mosquito assemblages near and among wetland habitats (*42, 43*). Traps were placed within the sedge habitat, but more than 60 m from the nearest focal flower patch to prevent any disturbance.

*P. obtusata* from the same site were enclosed in Bug Dorm cages (30cm x 30cm x 30cm; BioQuip® Products, Rancho Dominguez, CA, USA, # 1452) for which the bottom panel were removed to cover the orchid. Thirty mosquitoes were introduced into each cage through a sleeve located on the front panel and left without human interference for a duration of 48 h, after which the mosquitoes were collected from the enclosures and identified. The number and species of mosquitoes with pollinium attached were recorded, and plant was bagged for determination of the fruit-to-flower ratio at the end of the field season. A total of nineteen enclosures were used; 11 enclosures with a single plant, and 8 enclosures with 2-3 plants.

#### Pollen limitation studies

To determine the importance of pollination and out-crossing on *P. obtusata* fruit set, plants were subject to four different experimental treatments during the June through July summer months. For two weeks, plants were either unbagged (n = 20 plants) or bagged to prevent pollinator visitation (n = 19 plants). Organza bags (Model B07735-1; Housweety, Causeway Bay, Hong-Kong) were used to prevent pollinators from visiting the flowers. In addition, we determined the importance of cross- and self-pollination for *P. obtusata*. For cross pollination, six pollinia were removed from two plants using a toothpick and gently brushed against the stigma of a neighboring plant (n = 11 plants). To examine the effects of self-pollination, six pollinia were removed from three flowers and gently brushed the flowers on the same plant (n = 9 plants). At the end of the field season, the number of flowers and the number of fruits produced per individual plants were recorded and the fruit-to-flower ratios were calculated. For comparing the fruit weights and the seed set for each treatment, up to four fruits from each individual of *P. obtusata* were collected. The weights were measured with a digital scale (Mettler Toledo, Columbus, OH, USA), and the number of viable seeds per fruit were counted using an epifluorescent microscope (60x magnification; Nikon Ti4000). Fruit weights and seed sets were compared using a Student’s *t*-test; fruit-flower ratios were compared using a Mann-Whitney Test.

### Gas Chromatography coupled with Electroantennogram Detection (GC-EADs)

Electroantennogram signals were filtered and amplified (100×; 0.1-500 Hz) using an A-M 1800 amplifier (Sequim, WA, USA) connected to a personal computer via a BNC-2090A analog-to-digital board (National Instruments, Austin, TX, USA) and digitized at 20 Hz using WinEDR software (Strathclyde Electrophysiology Software, Glasgow, UK). A Hum Bug noise eliminator (Quest Scientific, Vancouver, Canada) was used to decrease electrical noise. The antennal responses to peaks eluting from the GC were measured for each mosquito preparation and for each peak and mosquito species. Bioactive peaks were those that elicited strong EAD responses, corresponding to deflections beyond the average noise floor of the baseline EAD signal. Responses by each individual preparation were used for Principal Component Analysis (Ade4 package, R). The responses of eight different mosquito species were tested to the scent extracts of three orchid species (n = 8 mosquito species for *P. obtusata*; n = 4 mosquito species each for *P. stricta* and *P. huronensis;* with 3-17 replicates per mosquito species per orchid, for a total of 109 GC-EAD experiments).

### Two-photon excitation microscopy

#### Calcium imaging in the *Ae. increpitus* mosquito AL

Odor-evoked responses in the *Ae. increpitus* mosquito antennal lobe (AL) with nine female mosquitoes at the beginning of the season when mosquitoes were relatively young (as defined by wing and scale appearance). Calcium imaging experiments were conducted using application of the calcium indicator Fluo4 to the mosquito brain and using a stage that allows simultaneous calcium imaging and tethered flight (*23*). The mosquito was cooled on ice and transferred to a Peltier-cooled holder that allows the mosquito head to be fixed to the stage using ultraviolet glue. The custom stage permits the superfusion of saline to the head capsule and space for movement by the wings and proboscis (*23*) (Fig. 3). Once the mosquito was fixed to the stage, a window in its head was cut to expose the brain, and the brain was continuously superfused with physiological saline (*22, 23*). Next, the perineural sheath was gently removed from the AL using fine forceps and 75 µL of the Fluo4 solution – made by 50 mg of Fluo4 in 30 µL Pluronic F-127 and then subsequently diluted in 950 µL of mosquito physiological saline – was pipetted to the holder allowing the brain to be completely immersed in the dye. Mosquitoes were kept in the dark at 15° C for 1.5 h (the appropriate time for adequate penetration of the dye into the tissue), after which the brain was washed 3 times with physiological saline. After the rinse, mosquitoes were kept in the dark at room temperature for approximately 10-20 min. before imaging.

Wing stroke amplitudes was acquired and analyzed using a custom camera-based computer vision system at frame rates of 100 Hz (*23, 44*), where the mosquito was illuminated with infrared LEDs (880 nm) and images were collected with an infrared-sensitive camera synched to the two-photon system. Stimulus-evoked initiation of flight and changes in the amplitude of the wing-stroke envelope were characterized for each odor stimulus (*sensu 23*). Calcium-evoked responses in the AL were imaged using the Prairie Ultima IV two-photon excitation microscope (Prairie Technologies) and Ti-Sapphire laser (Chameleon Ultra; Coherent). Experiments were performed at a depth of 40 µm from the ventral surface of the AL, allowing the calcium dynamics from approximately 18-22 glomeruli to be repeatedly imaged across preparations. Images were collected at 2 Hz, and for each odor stimulus images were acquired for 35 s, starting 10 s before the stimulus onset. Imaging data were extracted in Fiji/ImageJ and imported into Matlab (v2017; Mathworks, Natick, Massachusetts) for Gaussian filtering (2×2 pixel; σ = 1.5-3) and alignment using a single frame as the reference at a given imaging depth and subsequently registered to every frame to within ¼ pixel. Trigger-averaged ΔF/F were used for comparing glomerular responses between odor stimuli. After an experiment, the AL was sequentially scanned at 1 µm depths from the ventral to dorsal surface. Ventral glomeruli to the 40 µm depth were 3D reconstructed using Reconstruct software or Amira v5 (Indeed-Visual Concepts, Houston TX, USA) to provide glomerular assignment and registration between preparations. Glomeruli in the ventral region of the AL, based on their positions, were tentatively assigned names similar to those in *Ae. aegypti* (*23, 45*).

#### Calcium imaging in the *Ae. aegypti* mosquito AL

Odor-evoked responses in the *Ae. aegypti* AL were imaged taking advantage of our genetically-encoded *PUb-GCaMPs* mosquito line (*21*). A total of twenty preparations were used: 10 for single odorant and orchid mixture experiments; 6 for ratio experiments; and 4 for experiments using GABA-receptor antagonists. Glomeruli were imaged at 40 µm from the ventral surface, as glomeruli at this depth show strong responses to odorants in the orchid headspace, including nonanal, octanal, and lilac aldehyde, and at this depth approximately 14-18 glomeruli can be neuroanatomically identified and registered between preparations. Expression of GCaMP occurred in glia, local interneurons, and projection neurons. Nevertheless, double-labelling for GFP (GCaMPs) and glutamine synthase (GS; glial marker) revealed broad GFP labelling that did not always overlap with the glial stain, with GS-staining often occurring on astroglial-like processes on the rind around glomeruli, and strong GFP occurring within the glomeruli (Fig. S9). Thus, in our calcium imaging experiments we took care to image from the central regions of the glomeruli and avoid the sheaths and external glomerular loci. Moreover, strong GFP staining occurred in soma membranes located in the medial and lateral cell clusters, which contain the projection neurons and GABAergic local interneurons, respectively; the vast majority of these cell bodies did not stain for GS (Fig. S9). Relatedly, GCaMP6s expression is very high in AL local interneurons and projection neurons (PNs), such that during odor stimulation the PNs and axonal processes can often be imaged, and 3D reconstructions can be take place through simultaneous optical sections with odor stimulation. Nonetheless, we assume the glomerular responses are a function of multiple cell types. In other insects, GABAergic modulation has been shown to operate on olfactory receptor neurons, local interneurons and PNs (*28–30*).

Similar to experiments with *Ae. increpitus*, the majority the mosquitoes were UV-glued to the stage to allow free movement of their wings and proboscis; however, for experiments using GABA-receptor antagonists the proboscis was glued to the stage for additional stability. Once the mosquito was fixed to the stage, a window in its head was cut to expose the brain, and the brain was continuously superfused with physiological saline (*22*).

## Acknowledgments

We are grateful for the advice and assistance provided by G. Thornton, J. Patt, B. Nguyen, E. Lutz, J. Lim, E. Mathis, M. Clifford, and K. Moosavi. Photo images of *Platanthera* orchids and mosquitoes are courtesy of G. van Velsir, R. Coleman, T. Nelson, A. Jewiss-Gaines and F. Hunter. Funding: Support for this project was funded by National Institutes of Health under grants RO1-DC013693 (J.A.R.) and R21-AI137947 (J.A.R.); Air Force Office of Scientific Research under grants FA9550-14-1-0398 (J.A.R.) and FA9550-16-1-0167 (J.A.R.); an Endowed Professorship for Excellence in Biology (J.A.R.), and the University of Washington Innovation Award (J.A.R.).

## Supplementary Methods

### Floral VOCs collection and analysis

#### Orchid species

To characterize the orchid scents, headspace collections were performed during the summers of 2014, 2015 and 2016 in the Okanogan-Wenatchee National Forest (Washington, USA) and Yosemite National Park (California, USA). The scents of six *Platanthera* orchid species were studied: *P. obtusata* ((Banks ex Pursh) Lindley), the blunt-leaved orchid; *P. stricta* (Lindley), the slender bog orchid; *P. huronensis* (Lindley), the green bog orchid; *P. dilatata* (Pursh), the white bog orchid; *P. yosemitensis* (Colwell, Sheviak and Moore), the Yosemite bog orchid and *P. ciliaris* (Lindley), the yellow fringed orchid (Table S1). In the field, the plants were identified using a key (*1*). *P. ciliaris* was obtained from a nursery (Great Lakes Orchid LLC, Belleville, Michigan, USA) and maintained in the greenhouse of the Biology Department, at the University of Washington in Seattle, USA. Specimens of *P. obtusata*, *P. stricta* and *P. dilatata* were also maintained in the greenhouse as well as sampled in the field. For all orchid species, scents were collected during their peak flowering time and from those with unpollinated flowers.

#### Floral scent collection

To collect the flower scent, the inflorescence of the plant was enclosed in a nylon oven bag (Reynolds Kitchens, USA) that was tight around the stem. Two tygon tubes (Cole-Parmer, USA) were inserted at the base of the bag; one providing air into the bag through a charcoal filter cartridge (1 L/min.) to remove any contaminants from the pump or the surrounding air, and the other tube pulling the air out of the bag (1 L/min.) through a headspace trap composed of a borosilicate Pasteur pipette (VWR, Radnor, PA, USA) containing 50 mg of Porapak powder Q 80-100 mesh (Waters Corporation, Milford, MA, USA). This amount of Porapak was calibrated for collecting the orchid headspace without bleed through. The tubes were connected to a diaphragm pump (Diaphragm pump 400-1901, Barnant Co., Barrington, IL, USA for the greenhouse VOCs collection; Diaphragm pump 10D1125-101-1052, Gast, Benton Harbor, MI, USA, for the field VOCs collection connected to a Power-Sonic PS-6200 Battery, M&B’s Battery Company). Immediately after headspace collection, traps were eluted with 600 μL of 99% purity hexane (Sigma Aldrich, Saint-Louis, MO, USA). The samples were sealed and stored in 2 mL amber borosilicate vials (VWR, Radnor, PA) with Teflon-lined caps (VWR, Radnor, PA) on ice until reaching the laboratory, where they were stored at −80°C until analysis by GCMS. Because some orchid species are pollinated by nocturnal moths (e.g., *P. dilatata*), whereas others are pollinated by diurnal insects (e.g., *P. obtusata*), we elected to normalize collections across *Platanthera* species for an entire 24 h period, excepting those of *P. ciliaris* which was collected for 72 h to account for the chemical analyses and relative abundance in the scent. For headspace controls, samples were taken concurrently from empty oven bags and from the leaves of the plants (as vegetation-only controls). 7-39 floral headspace collections were conducted for each orchid species for a total of 109 floral headspace samples.

#### Gas Chromatography with Mass Spectrometric Detection of the orchid scents

One to three microliters of each sample were injected into an Agilent 7890A GC and a 5975C Network Mass Selective Detector (Agilent Technologies, Palo Alto, CA, USA). A DB-5 GC column (J&W Scientific, Folsom, CA, USA; 30 m, 0.25 mm, 0.25 μm) was used, and helium was used as the carrier gas at a constant flow of 1 cc/min. For runs with the DB-5 column, the oven temperature was 45° for 4 min, followed by a heating gradient of 10° to 230°, which was then held isothermally for 6 min. The total run time was 28.5 min. A Cyclosil-B column (J&W Scientific, Folsom, CA, USA; 30 m, 0.25 mm, 0.25 μm) was used to examine the stereoisomer composition of the lilac aldehyde in the floral scents. For the chiral column, the oven temperature was 45° for 6 min, followed by a heating gradient of 5° to 160°, then 15° to 200° which was then held isothermally for 5 min. The total run time was 36.7 min. Natural lilac aldehydes were isolated from lilac flowers (*Syringa vulgaris*) to create a natural standard, because lilac flowers are known to contain 4 out of 8 possible lilac aldehyde stereoisomers, all of which have the 5’S configuration. The natural standard was prepared by purifying the lilac aldehydes from *Syringa vulgaris* flowers by CO_2_ extract (Hermitage Oils, Petrognano, IT) using column chromatography with Silica Gel 60, mesh 230 - 400 (Material Harvest Ltd, Cambridge, UK), and eluted with 90% hexanes, 10% ethyl acetate. 1 µl of the sample was injected into the GCMS with the chiral column. The lilac aldehyde peaks from *Platanthera* samples were matched with peaks from lilac aldehyde purified from lilac flower CO_2_ extract using the ChemStation software (Agilent Technologies, Santa Clara, CA, USA). The lilac aldehyde peaks in the samples, and in the standard purified from lilac flower CO_2_ extract were matched based on their retention indices.

Chromatogram peaks were then manually integrated using the ChemStation software (Agilent Technologies, Santa Clara, CA, USA) and tentatively identified by the online NIST library. Using methods developed in our laboratory for identifying and quantifying volatiles in floral headspace emissions (*2–4*), the data from each sample was first run through a custom program (https://github.com/cliffmar/GCMS_and_combine) to identify the volatiles based on their Kovats index and to remove potential contaminants and chemical synonyms for the subsequent analyses.

Synthetic standards at different concentrations (0.5 ng/µl to 1 µg/µl) were then run to identify the peaks further and to quantify the areas for each compound; peaks are presented in terms of nanograms per hour per inflorescence (Table S3). Results were plotted and analyzed using a Non-metric multidimensional scaling (NMDS) analysis with a Wisconsin double standardization and square-root transformation of the emission rates and the Bray–Curtis dissimilarity index on the proportions using the *vegan* package in R (*5*). We then performed an ANOSIM on the data, allowing us to statistically examine differences in chemical composition and relative abundance between orchid species.

### Mosquitoes rearing and colony conditions

#### Field mosquitoes

Adult mosquitoes were caught by hand, using plastic containers (BioQuip® Products, Rancho Dominguez, CA, USA), on the sites where the orchids were located. We also collected pupae in ponds located in the same areas. The mosquitoes were then brought back to the lab, maintained in cages (BioQuip® Products, Rancho Dominguez, CA, USA) and placed in environmental chambers (22±1°C during the day and 17±1°C during the night, 60±10% relative humidity (RH) and with a 12-12 h light-dark cycle). There, they had access to 10% sucrose *ad libitum*. Before the experiments, the mosquitoes were starved for two days, CO_2_ anesthetized (Flystuff Flypad, San Diego, CA, USA) and identified using standard keys (*6, 7*). We used the taxonomic naming convention of Wilkerson et al. (2015) for classifying the field-caught mosquitoes (*8*). The mosquitoes bearing pollinia were snap frozen in liquid nitrogen for further analyses.

#### Laboratory mosquito strains

Female *Ae. aegypti* (wild type, MRA-734, ATCC®, Manassas, VA, USA) and *An. stephensi* (MRA-128, Strain STE2, CDC, Atlanta, GA, USA) mosquitoes were also used for behavioral experiments. Mosquitoes were kept in an environmental chamber maintained at 25 ± 1°C, 60 ± 10% RH and under a 12-12 h light-dark cycle. Groups of 200 larvae were placed in 26×35×4cm covered trays containing tap water and were fed daily on fish food (Hikari Tropic 382 First Bites - Petco, San Diego, CA, USA). Groups of same age pupae (both males and females) were then isolated in 16 oz containers (Mosquito Breeder Jar, BioQuip® Products, Rancho Dominguez, CA, USA) until emergence. Adults were then transferred into mating cages (BioQuip® Products, Rancho Dominguez, CA, USA) and maintained on 10% sucrose. An artificial feeder (D.E. Lillie Glassblowers, Atlanta, Georgia, USA; 2.5 cm internal diameter) filled with heparinized bovine blood (Lampire Biological Laboratories, Pipersville, PA, USA) placed on the top of the cage was heated at 37°C using a water-bath circulation system (HAAKE A10 and SC100, Thermo Scientific, Waltham, MA, USA) and used to feed mosquitoes weekly. For the experiments, groups of 120 pupae were isolated and maintained in their container for 6 days after their emergence. Mosquitoes had access to 10% sucrose but were not blood-fed before the experiments. The day the experiments were conducted, mosquitoes were cold-anesthetized (using a climatic chamber at 10°C) and females were selected manually with forceps.

*Ae. aegypti PUb-GCaMP6s* mosquitoes (*9*) used in the calcium imaging experiments were from the Liverpool strain, which was the source strain for the reference genome sequence. Briefly, this mosquito line was generated by injecting a construct that included the GCaMP6s plasmid (ID# 106868) cloned into the piggyBac plasmid pBac-3xP3-dsRed and using *Ae. aegypti* polyubiquitin (*PUb*) promoter fragment. Mosquito pre-blastoderm stage embryos were injected with a mixture of the GCaMP6s plasmid described above (200ng/ul) and a source of piggyBac transposase (phsp-Pbac, (200ng/ul)). Injected embryos were hatched in deoxygenated water and surviving adults were placed into cages and screened for expected fluorescent markers. Mosquitoes were backcrossed for five generations to our wild-type stock, and subsequently screened and selected for at least 20 generations to obtain a near homozygous line. The location and orientation of the insertion site was confirmed by PCR (see (*9*) for details).

All behavioral and physiological experiments were conducted at times when mosquitoes were the most active (*10, 11*).

### Preparation for Gas Chromatography coupled with Electroantennogram Detection (GC-EADs)

Individual mosquitoes were isolated in falcon^TM^ tubes (Thermo Fisher Scientific, Pittsburgh, PA, USA) covered with a piece of fine mesh. They were maintained in a climatic chamber, as previously described, and identified immediately before running the experiment. Carbon dioxide delivered through a pad (Genesee Scientific, San Diego, CA, USA) was used to briefly anesthetize mosquitoes before transferring them on ice for the dissection. The head was excised and the tip (i.e., one segment) of each antenna was cut off with fine scissors under a binocular microscope (Carl Zeiss, Oberkochen, Germany). The head was then mounted on an electrode composed of a silver wire 0.01” (A-M Systems, Carlsbord, WA, USA) and a borosilicate pulled capillary (Sutter Instrument Company, Novato, CA, USA) filled with a 1:3 mix of saline^42^ and electrode gel (Parker Laboratories, Fairfield, NJ, USA) in order to avoid the preparation to desiccate during the experiment. The head was mounted by the neck on the reference electrode. The preparation was then moved to the GC-EAD with the tips of the antennae inserted under the microscope (Optiphot-2, Nikon, Tokyo, Japan) into a recording electrode, that was identical to the reference electrode. The mounted antennae were oriented at 90° from the main airline which was carrying filtered air (Praxair, Danbury, CT, USA) and volatiles eluting from the Gas-Chromatograph to the preparation via a 200° C transfer line (EC-05; Syntech GmbH, Buchenbach, Germany). Five microliters of the orchid extract was injected into the Gas Chromatograph with Flame Ionization Detection (Agilent 7820A GC, Agilent Technologies; DB5 column, J&W Scientific, Folsom, CA, USA). The oven program was the same as the one used for the GC-MS analyses of the scent extracts. The transfer line from the GC to the preparation was set to 200° C.

### Behavioral experiments

#### Chemical mixture preparation and single odorants

All the chemicals used for the behavioral experiments were ordered from Sigma Aldrich (St. Louis, MO, USA)(≥98% purity) with the exception of the lilac aldehyde (mixture of B [49%], D [26%], and C [23%] isomers) that were synthesized by Medchem Source LLP (Seattle, WA, USA) according to the methods of Wilkins et. al. (*52*). The ratio of D and C isomers approximated those quantified in the *P. obtusata* scent (Table S3). Briefly, linalool (0.5 mL) was aliquoted in dioxane (2 mL) and subsequently stirred with selenium dioxide (225 mg) under reflux for approximately 6 h. Afterward the solution was separated using silica gel yielding 5-dimethyl-5-ethenyl-2-tetrahydrofuranacetaldehydes (lilac aldehyde, mixture of isomers). Purity was verified by two-dimensional COSY NMR (AC-300, Bruker, Billerica, MA) and GC-MS (Agilent Technologies, Palo Alto, CA, USA).

Stimuli included the scent from live *P. obtusata* flowers; artificial mixture of the *P. obtusata* scent (with or without the lilac aldehyde); the lilac aldehyde at the concentration in the *P. obtusata* scent mixture; and the negative mineral oil (no odor) control. The artificial mixture was composed of a 14 component blend of odorants identified as antennal-active (*via* the GC-EAD experiments)(Table S3): The mixture was prepared by adding each synthetic component and adjusting so that the headspace concentrations matched those found in the *P. obtusata* floral headspace (as quantified through GC-MS). Briefly, emission rates of the artificial mixtures and single compounds were scaled to those of live flowers by their individual vapor pressures and associated partial pressures, and verified and adjusted by iterative headspace collection and quantification using the GC-MS (*sensu* (*3, 4*)).

To test the effects of different ratios of lilac aldehyde in the orchid scents, mixtures were created where the composition and concentration of volatiles were the same as those in the *P. obtusata* scent, except we increased the concentration of the lilac aldehyde to similar levels as those measured in the scents of *P. stricta*, *P. dilatata*, and *P. huronensis* (Table S3). Finally, higher tested concentrations of the *P. obtusata* mixture – well beyond those emitted naturally by *P. obtusata* plants – were significantly aversive to the mosquitoes (binomial tests: p < 0.05).

#### Olfactometer

Female *Ae. aegypti* (MRA-734; n = 645 tested and flew; n = 482 made a choice) and *An. stephensi* (MRA-128; n = 153 tested and flew; n = 73 made a choice) from our laboratory colonies, and *Ae. increpitus* and *Ae. communis* caught in the field (n = 138 tested), were used for these experiments. Female mosquitoes were individually selected and checked for the integrity of their legs and wings to ensure that they would be able to behave properly in the olfactometer. Females were then individually placed in 50 mL conical Falcon^TM^ tubes (Thermo Fisher Scientific, Pittsburgh, PA, USA) covered by a piece of mesh maintained by a rubber band.

A custom-made Y-maze olfactometer made from Plexiglas® was used to compare behavioral response of the mosquitoes to different odor stimuli. The olfactometer is comprised of a starting chamber where the mosquitoes were released, a tube (length: 30 cm; diameter 10 cm) connected to a central box where two choice arms of equal length (39 cm) and diameter (10 cm) were attached. Fans (Rosewill, Los Angeles, CA, USA) placed inside a Plexiglas® box were connected to the two arms of the olfactometer. The fans generate airflows of 20 cm/s. The air first passes through a charcoal filter (C16×48, Complete Filtration Services, Greenville, NC, USA) to remove any odor contaminants that may be in the ambient air. The filtered air then passes through a mesh screen and an aluminum honeycomb core (10 cm in thickness) to create a laminar flow within the olfactometer. Odor delivery to each choice arm is made using an aquarium pump adjusted with flow meters (Cole-Parmer, Vernon Hills, IL, USA). Air lines (Teflon® tubing; 3 mm internal diameter) were connected to one of two 20 mL scintillation vials containing the odor stimulus or control (mineral oil). Odor stimuli were deposited on Whatman® Grade 1 Filter Paper (32 mm diameter, VWR International, Radnor, PA USA) cut into strips (1 cm x 5 cm). Each line was connected to the corresponding choice arm of the olfactometer and placed at about 4 cm from the fans so that the tip of the tubing was centered in the air flow generated by the fans, and flow through the tubes was approximately 5 mL/min. To ensure the odor stimuli did not decrease in concentration over the course of the experiment, the odor-laden filter papers were replaced every 20 to 25 minutes and verified by SPME and GCMS. All the olfactometer experiments were conducted in a well-ventilated environmental chamber (Environmental Structures, Colorado Springs, CO, USA) maintained at 25°C and 50-70% RH. After each experiment, the olfactometer, tubing and vials were sequentially cleaned with 70% and 95% ethanol and dried overnight to avoid any contamination between experiments. Finally, to control for any directional biases, the control- and odor-bearing arms of the olfactometer were randomized between experiments. A Logitech C615 webcam (Logitech® Newark, CA, USA) mounted on a tripod and placed above the olfactometer was used to record the mosquito activity during the entire experiment.

The experiment began when one single mosquito was placed in the starting chamber. The mosquito then flew along the entry tube and, at the central chamber, could choose to enter one of the olfactometer arms, one emitting the odor and the other the “clean air” (solvent only) control (*11*). We considered the first choice made by mosquitoes when they crossed the entry of an arm. Mosquitoes that did not choose or did not leave the starting chamber were considered as not responsive and discarded from the preference analyses. In addition, to ensure that contamination did not occur in the olfactometer and to test mosquitos’ responses to innately attractive, mosquitoes were placed in the olfactometer and exposed to either two clean air currents (neutral control). Overall, approximately 60% of the females were motivated to leave the starting chamber of the olfactometer and choose between the two choice arms.

Binary data collected in the olfactometer were analyzed and all statistical tests were computed using R software (R Development Core Team (*13*)). Comparisons were performed by means of the Exact Binomial test (α=0.05). For each treatment, the choice of the mosquitoes in the olfactometer was either compared to a random distribution of 50% on each arm of the maze or to the distribution of the corresponding control when appropriate. For binary data, the standard errors (SE) were calculated as in (*11*):

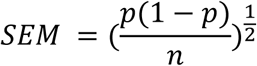

For each experimental group, a preference index (PI) was computed in the following way: PI = [(number of females in the test arm – number of females in the control arm) / (number of females in the control arm + number of females in the test arm)]. A PI of +1 indicates that all the mosquitoes chose the test arm, a PI of 0 indicates that 50% of insects chose the test arm and 50% the control arm, and a PI of −1 indicates that all insects chose the control arm of the olfactometer (*11*).

### Two photon experiments

#### Glomerular registration from two photon experiments

We initially attempted to register glomeruli using a previously published AL atlas (*14*), but the number, position and size of glomeruli from our imaging experiments did not always match those of the previous study. We thus created a provisional atlas with female mosquitoes (n = 6) that allowed us to cross-reference the imaged glomeruli and compare their positions and structures to those described in (*14*). This was accomplished via clear glomerular boundaries, especially during odor stimulation, and the distinct odorant tuning of glomeruli throughout the depths of the AL (e.g., AM2 responsive to DEET; LC2 and AL3 responsive to nonanal; PD3 responsive to geosmin; and MD2 responsive to CO_2_) that allowed 3D registration across preparations and subsequent warping and referencing with the atlas. Based on these results we tentatively assigned glomerular names. Nonetheless, future work will be needed to enable the olfactory receptor inputs to their cognate glomeruli. Fortunately, thanks to the recent development of new genetic tools (*9, 15, 16*), these types of experiments will soon be possible.

#### Saline and pharmacological agents

The saline was made based on the Beyenbach and Masia recipe (*17*) and contained 150.0 mM NaCl, 25.0 mM N-2-hydroxyethyl-piperazine-N’-2-ethanesulfonic acid (HEPES), 5.0 mM sucrose, 3.4 mM KCl, 1.8 mM NaHCO_3_, 1.7 mM CaCl_2_, and 1.0 mM MgCl_2_. The pH was adjusted to 7 with 1 M NaOH. Immediately before the experiment, GABA receptor antagonists were dissolved in saline (1 µM Picrotoxin (Sigma-Aldrich, St. Louis, MO; P1675), and 10 µM CGP54626 (Tocris Bioscience, Park Ellisville, MO; CGP 54626); to block both GABA-A and GABA-B receptors). A drip system comprising two 100 mL reservoirs – one containing the GABA receptor antagonists, and the other saline – converged on the two-channel temperature controller to facilitate rapid switching from normal physiological saline solution to the antagonists and back. Antagonists were superfused directly into the holder near to the opening of the head capsule and recorded neuropil. The odor-evoked responses were first recorded under normal physiological saline solution and then repeated under GABA receptor antagonists diluted in normal saline solution, and finally the normal saline wash. All calcium imaging data were statistically analyzed using Kruskal-Wallis tests with multiple comparisons and visualized using Principal Components Analysis. Analyses were performed in Matlab (v2017; Mathworks, Natick, Massachusetts).

#### Olfactory delivery and stimuli

Olfactory stimuli were delivered to the mosquito by pulses of air diverted through a 2 mL cartridge containing a piece of filter paper bearing the odor stimulus (2 µL). An airline provided gentle and constant charcoal-filtered air at 1 m/s to the antennae allowing continuous ventilation to prevent adaptation of the olfactory receptor cells. The stimulus output was positioned in the airline 2 cm from and orthogonal to the mosquito antennae. For testing different ratios of lilac aldehyde and nonanal, two syringes bearing different concentrations of the odorants were used and positioned such that the outputs were positioned in the same location in the air stream. Pulses of odor, each at a duration of two seconds and at a flow rate of approximately 5 ml/min., were delivered to the antennae using a solenoid-activated valve (The Lee Company, Essex, CT, USA, LHDA0533115H) controlled by the PrairieView software. Odor stimuli were separated by intervals of 120 s to avoid receptor adaptation. The two-way valve enabled a constant airstream with minimal disturbance during odor stimulation. Odorants (>98% purity; Sigma-Aldrich, St. Louis, MO, USA) were diluted in mineral oil to scale the intensities to those quantified in the *P. obtusata* scent, except for DEET (N,N-diethyl-meta-toluamide)(1-40% concentrations) which was diluted in 200 proof ethanol, and each cartridge used for no more than 4 stimulations. Olfactory stimuli were: aliphatics (nonanal [220 ng], octanal 46 ng], hexanal [9 ng], 1-octanol [0.5 ng]); monoterpenes (α-pinene [1.44 ng], β-pinene [1.5], camphene [1.41 ng], β-myrcene [3.5 ng], D-limonene [16.5 ng], eucalyptol [3.4 ng], lilac aldehyde (B, C and D isomers) [124.7 ng], (±)linalool [1.41 ng], and myrtenol [1.35 ng]); aromatics (benzaldehyde [1.45 ng], DEET [10%]); and mixtures, including human scent, the *P. obtusata* mixture, the *P. stricta* mixture, the *P. dilatata* mixture, and the *P. huronensis* mixture. Similar to behavioral experiments, for experiments examining the effects of lilac aldehyde in the flower mixtures (Fig. 4C,D), the odor constituents were kept the same except for the lilac aldehyde, which was scaled to the headspace concentrations of *P. strica*, *P. huronensis*, or *P. dilatata*. This provided a mechanism to determine how the change of one odorant concentration in the mixture impacted the activation or suppression of glomeruli in the ensemble. Importantly, all odorant constituents and floral mixtures were at the same headspace concentration levels as the natural floral scents or scent constituents, as verified by headspace collections and quantification using the GC-MS.

Human scent samples were collected by gently rubbing Whatman filter paper on the ankles and wrists of one human volunteer per experiment. Prior to human scent collection, volunteers placed their ankles and wrists over running water for ten minutes. The human scent protocols were reviewed and approved by the University of Washington Institutional Review Board, and all volunteers gave their informed consent to participate in the research. Control solvents for the olfactory stimuli were mineral oil (for the majority of odorants and mixtures) and ethanol (for DEET).

### Immunohistochemistry

To register putative glomeruli in our calcium imaging experiments, we created an AL atlas using antiserum against tyrosine hydroxylase (ImmunoStar, Hudson, WI, USA - Cat. no. 22941; 1:50 concentration), GABA (Sigma-Aldrich, St. Louis, MO, USA - Cat. no. A2052; 1:100 concentration) and monoclonal antisera against alpha-tubulin (12G10; 1:1000 concentration; developed by Drs. J. Frankel and E.M. Nelsen). In addition, to characterize the expression of GCaMP in different cell types in the AL, we also double-stained for GFP (for the GCaMP6s; Abcam, Cambridge, MA, USA – Cat. no. ab6556; 1:1000 concentration) and glutamine synthase (GS; a glial marker; Sigma-Aldrich, St. Louis, MO, USA - Cat. no. MAB302; 1:500 concentration); and GABA and GS. The alpha-tubulin antiserum was obtained from the Developmental Studies Hybridoma Bank developed under the auspices of the NICHD and maintained by the University of Iowa, Department of Biology (Iowa City, IA). These antisera either provide clear designation of glomerular boundaries, allowing 3D reconstruction of individual glomeruli, or designation of glial-, GABA-, and GFP-stained cells and processes. Briefly, animals were immobilized by refrigeration at 4° C and heads were removed into cold (4° C) fixative containing 4% paraformaldehyde in phosphate-buffered saline, pH 7.4 (PBS, Sigma-Aldrich, St. Louis, MO, USA-Cat. No. P4417). Heads were fixed for 1 h and then brains were dissected free in PBS containing 4% Triton X-100 (PBS-TX; Sigma-Aldrich, St. Louis, MO, USA - Cat. No. X100). Brains were incubated overnight at 4° C in 4% PBS-TX. Brains were washed three times over 10 minutes each in 0.5% PBS-TX and then embedded in agarose. The embedded tissue was cut into 60 µm serial sections using a Vibratome and washed in PBS containing 0.5% PBS-TX six times over 20 minutes. Then 50 µL normal serum was added to each well containing 1,000 µL PBS-TX. After 1 hour, primary antibody was added to each well and the well plate was left on a shaker overnight at room temperature. The next day, sections were washed six times over 3 h in PBS-TX. Then 1,000-µL aliquots of PBS-TX were placed in tubes with 2.5 µL of secondary Alexa Fluor 488 or Alexa Fluor 546-conjugated IgGs (ThermoFisher Scientific, Waltham, MA, USA) and centrifuged at 13,000 rpm for 15 minutes. A 900-µL aliquot of this solution was added to each well, and tissue sections were then washed in PBS six times over 3 h, embedded on glass slides in Vectashield (Vector Laboratories, Burlingame, CA, USA - Cat. No. H-1000) and imaged using a Leica SP5 laser scanning confocal microscope. Images were processed using ImageJ (National Institutes of Health) and a 3D atlas, assembled from 6 mosquitoes, were constructed using the Reconstruct software (v. 1.1.0.0)(*18*).

---

**Fig. S1.**
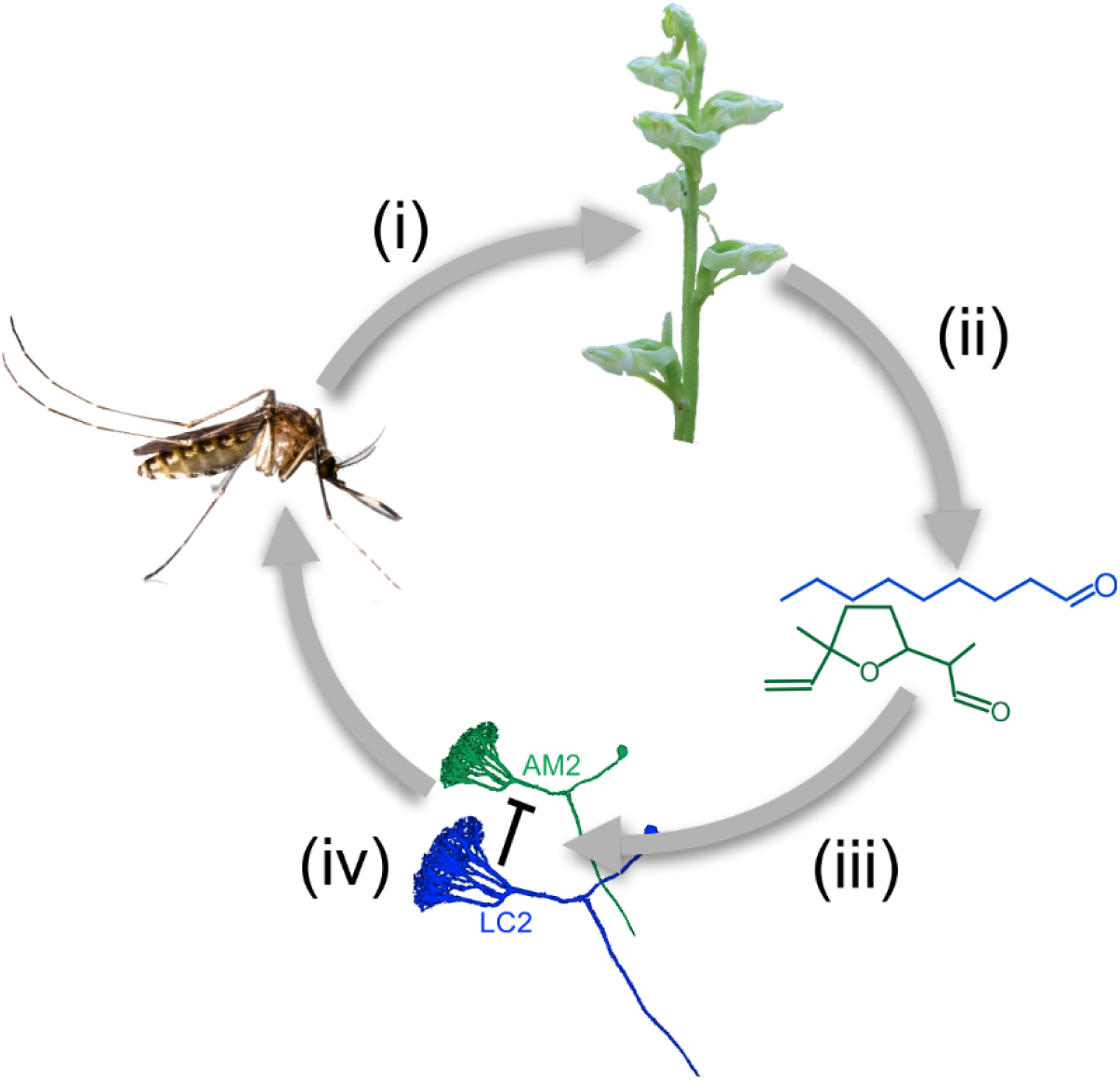
Experimental series for determining the olfactory basis of the *Aedes-Platanthera* mutualism. An integrative series of experiments were performed to evaluate the behavioral and neural bases of the *Aedes-Platanthera* relationship, including: (*i*) pollination experiments; (*ii*) chemical analytical studies of the orchid scents; (*iii*) identification of antennal responsive volatiles; and (*iv*) calcium imaging experiments in the mosquito AL to characterize the glomerular representations of the orchid scents. Together, these experiments allowed us to test the working hypothesis that for *Aedes* mosquitoes’ certain odorants play critical roles in the detection and discrimination of floral scents, and inhibition in the AL is essential for scent discrimination.

**Fig. S2.**
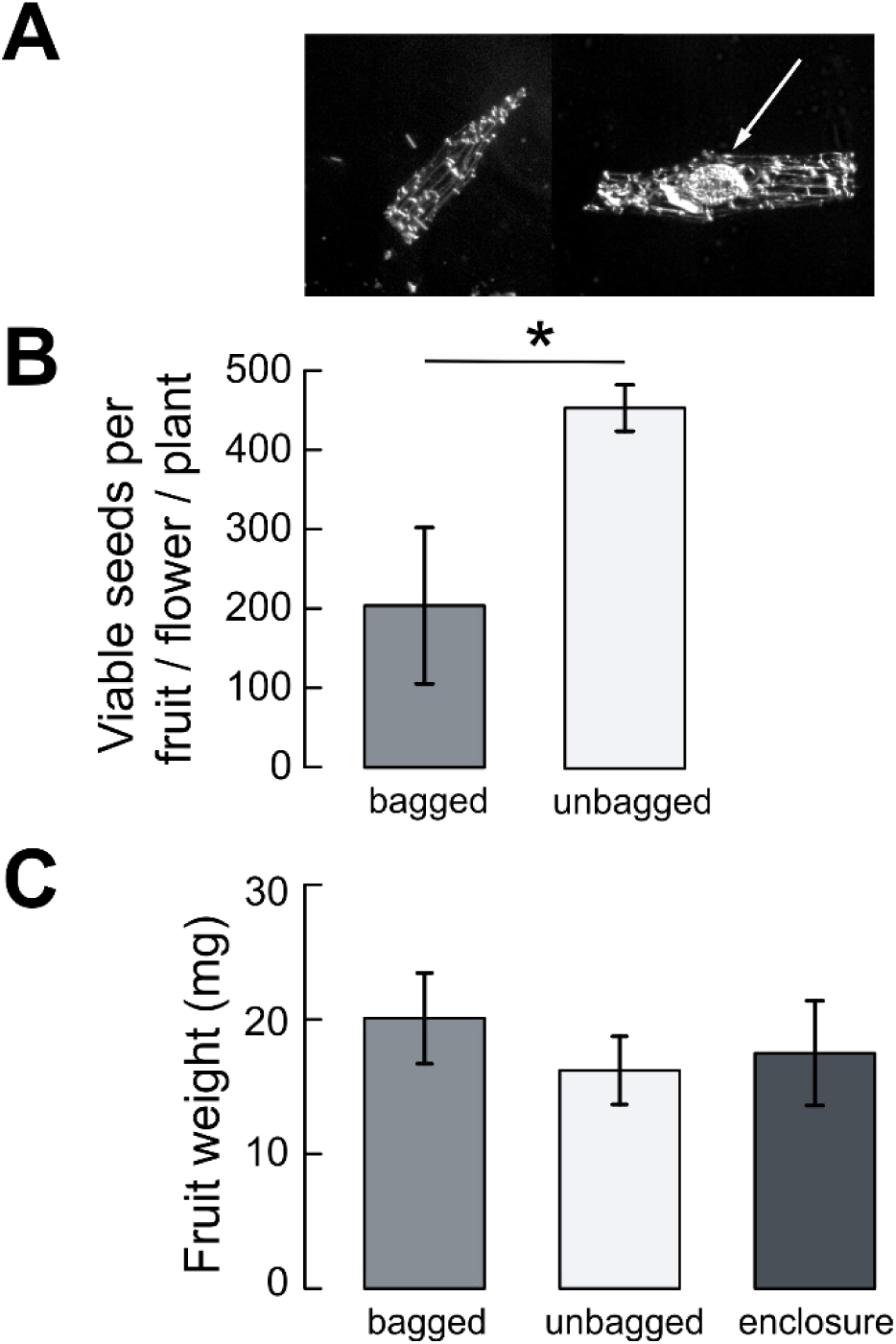
Seed set and fruit weight for *P. obtusata* pollen limitation and enclosure experiments. (**A**) *P. obtusata* fruits were opened and viable seeds identified by the identifying the embryo within the seed capsule (arrow). (**B**) The number of viable seeds per flower per plant for bagged and unbagged plants. Bars are the mean ± SEM; asterisk denotes significant difference between treatments (Student’s *t* test: p<0.05). (**C**) Fruit weights for plants in the unbagged, bagged, and enclosure treatments. Bars are the mean ± SEM; there was no significant different between treatments (Student’s *t* test: p>0.05).

**Fig. S3.**
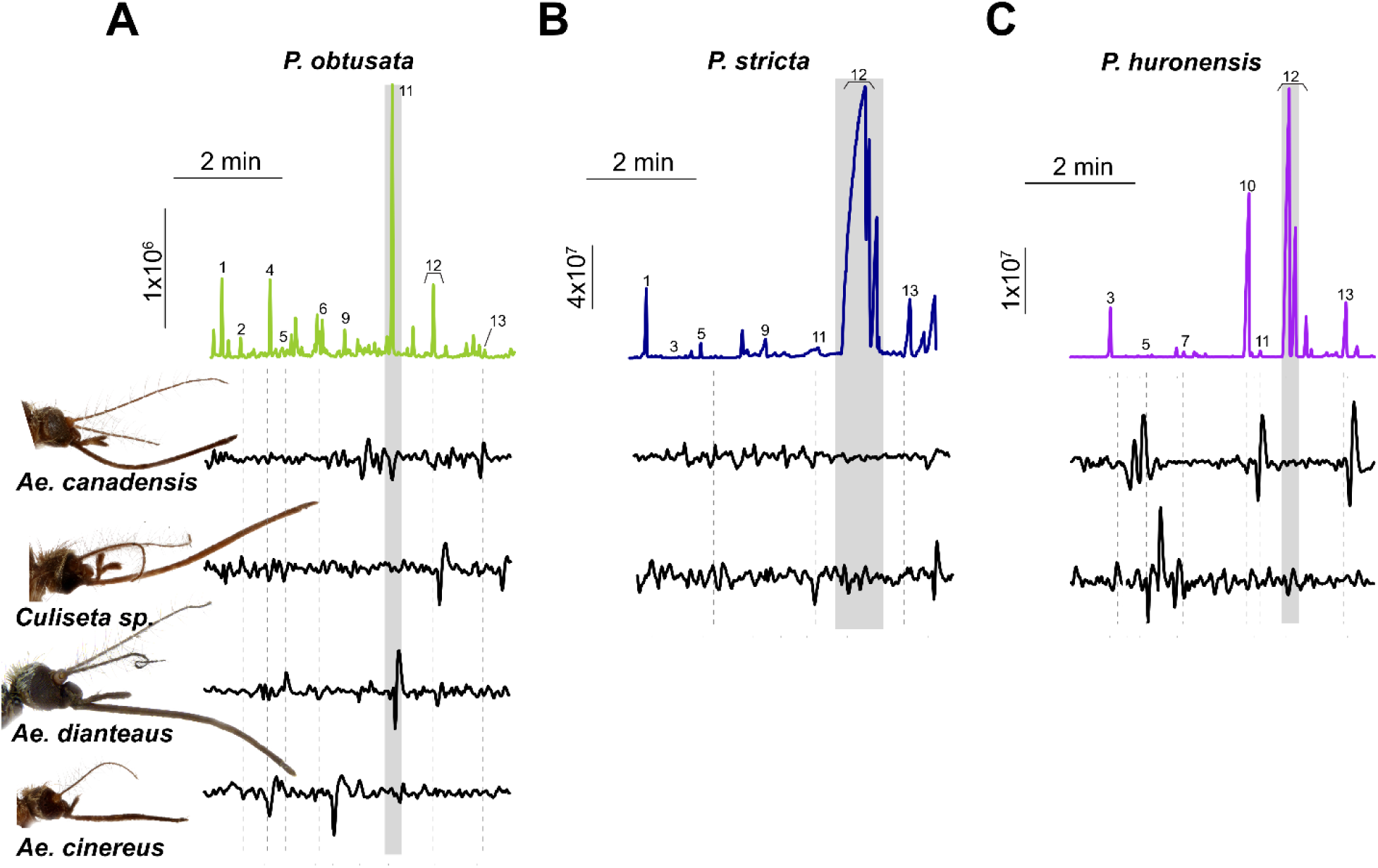
Identification of antennal responsive orchid volatiles in mosquitoes. As in Figure 2, Gas chromatogram traces for the *P. obtusata* (**A**), *P. stricta* (**B**), and *P. huronensis* (**C**) headspaces, with individual electroantennogram responses to the GC peaks for four mosquito groups (*Ae. canadensis*, *Culiseta sp., Ae. dianteaus*, and *Ae. cinereus*) below. (1) α-pinene, (2) camphene, (3) benzaldehyde, (4) β-pinene, (5) β-myrcene, (6) octanal, (7) D-limonene, (8) eucalyptol, (9) 1-octanol, (10) linalool, (11) nonanal, (12) lilac aldehyde (C, D isomers), (13) lilac alcohol. For each species, electroanntenogram responses from each individual mosquito are shown in Figure 2B (*n=*3-16 mosquitoes per species).

**Fig. S4.**
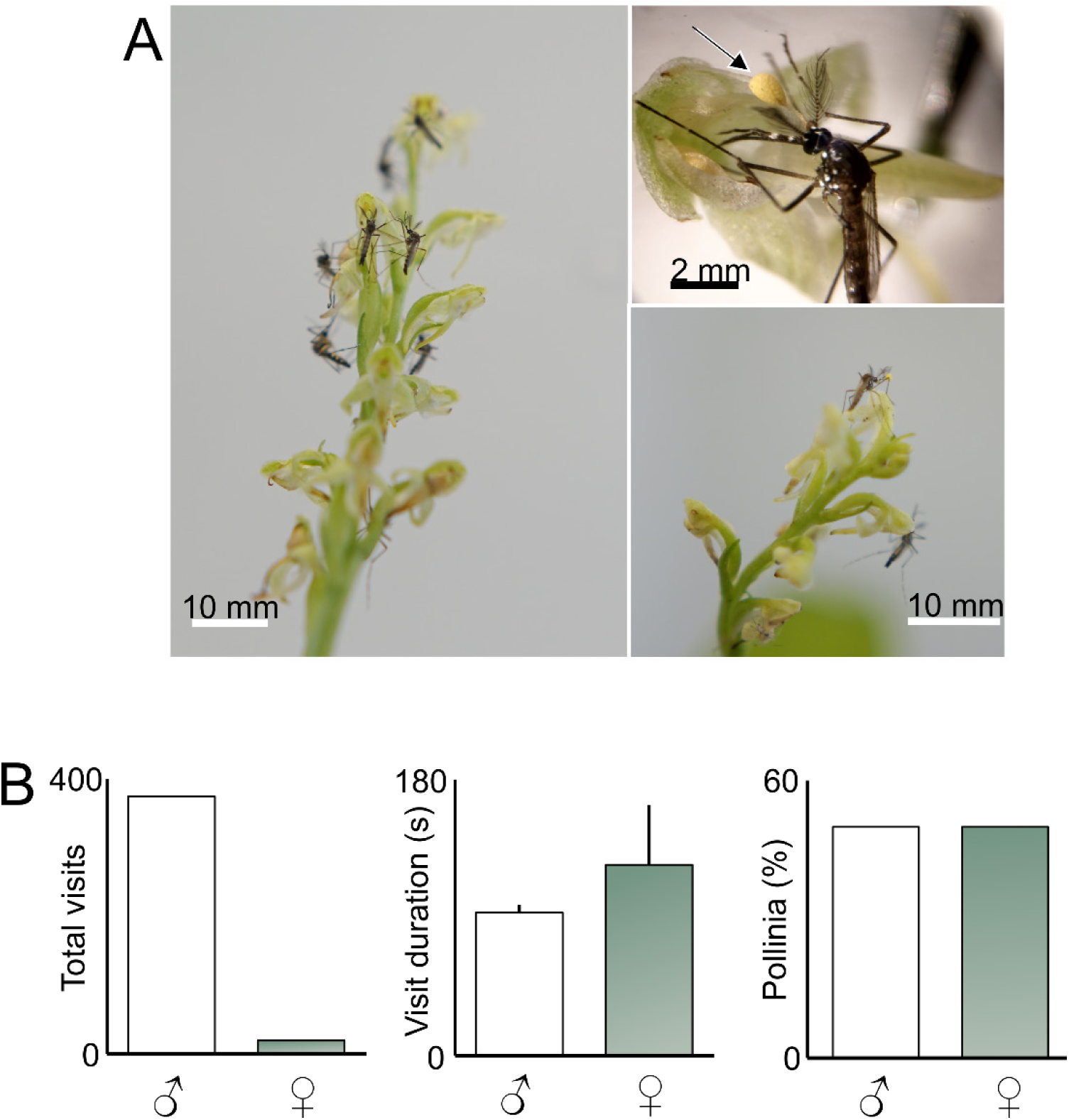
*Ae. aegypti* orchid visitation and pollinia attachment. 50 *Ae. aegypti* were released into enclosures containing a plant (n = 3; with 21 total flowers). Mosquito-orchid observations were taken by video or direct observations for approximately 5h. (**A**) Once released into the enclosure, both male and female *Ae. aegypti* mosquitoes landed and began probing the flowers and inserting their proboscis into the nectar spur. Similar to *Ae. communis* and *Ae. increpitus*, after insertion the pollinia would often be attached to the eye (arrow points to pollinia). (**B**) The total number of *Ae. aegypti* plant visits (left), visit duration (middle), and percentage of pollinia attachment for male and female mosquitoes. Although more male mosquitoes visited the *P. obtusata* plant, there was no statistical difference in the visit duration (*t-*test: p<0.05) or pollinia attachment between sexes (binomial test: p=0.5).

**Fig. S5.**
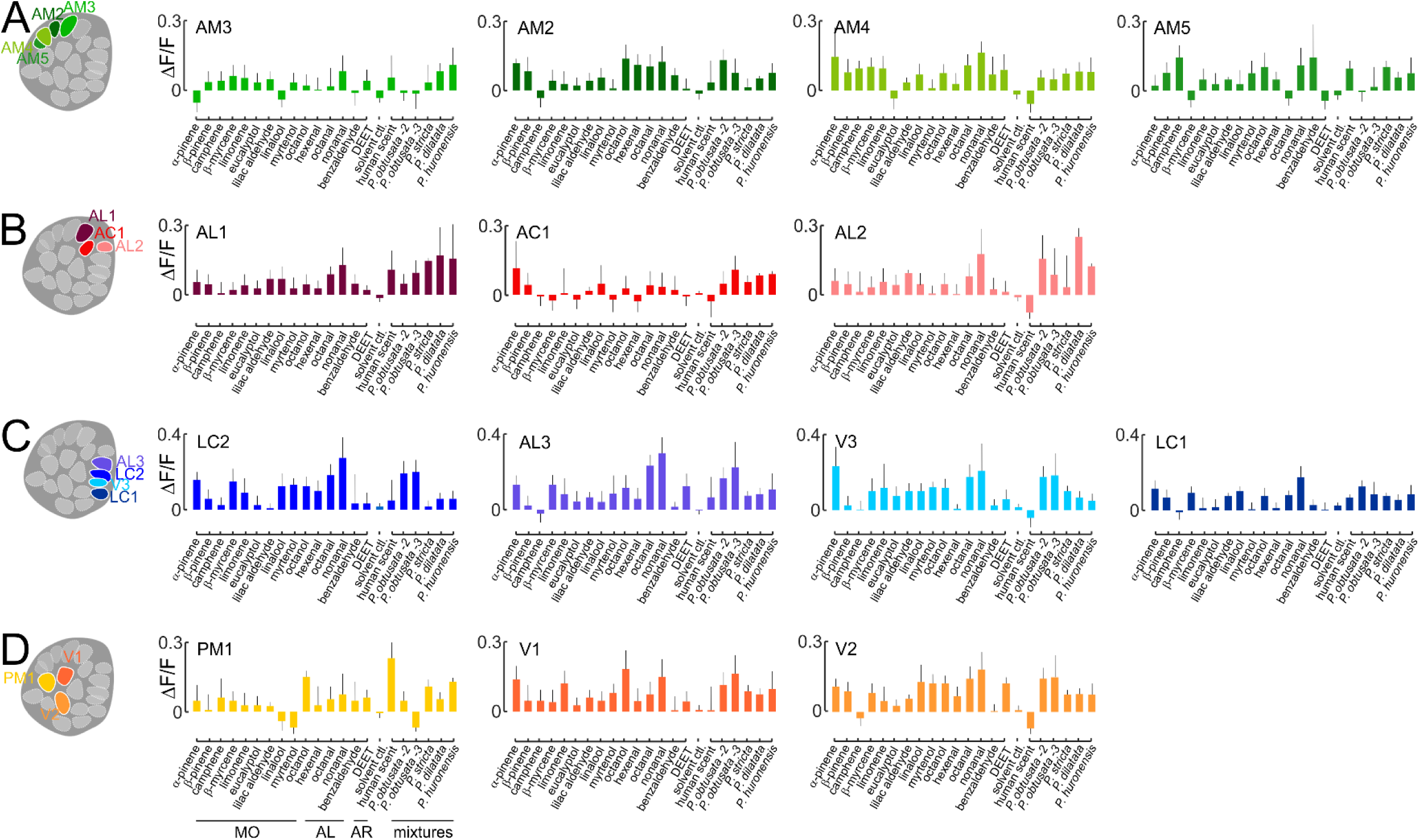
*Ae. increpitus* glomerular responses to odors. Using a ΔF/F threshold of 0.15, 14 of the 22 glomeruli imaged from the *Ae. increpitus* mosquito AL elicited strong responses to a panel of odorants identified in the *P. obtusata* headspace, DEET (another bioactive odorant), mixtures including the orchid scents and human scent, and the mineral oil (no odor) control. Odorants of the different chemical classes elicited distinct responses in glomeruli (Kruskal-Wallis test: p < 0.005), and glomerular clusters were significantly different in their responses (p < 0.001). (**A**) (Left) Location of the imaged glomeruli within the imaging plane. Glomeruli were assigned names similar to those of *Ae. aegypti* based on their position and morphology. (Right) Responses of the AM2, AM3, AM4 and AM5 glomeruli to odor stimuli. Bars are the mean ± SEM (n = 5-9 mosquitoes) (**B**) As in A, except for the AL1, AL2 and AC1 glomeruli. (**C**) As in A, except for the AL3, V3, LC1 and LC2 glomeruli. (**D**) As in A, except for the V1, V2 and PM1 glomeruli.

**Fig. S6.**
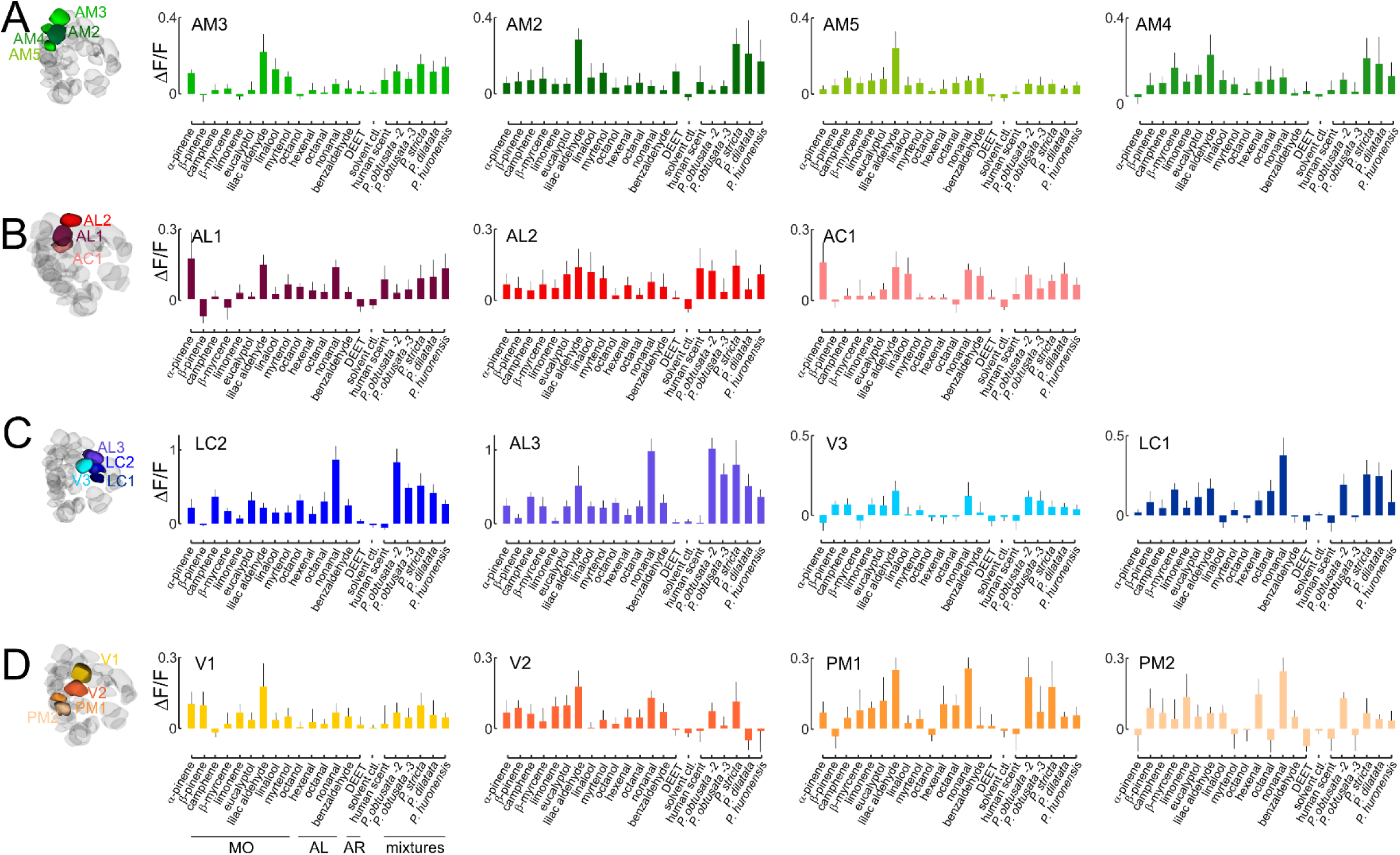
*Ae. aegypti* glomerular responses to odors. As in Figure S5: using a ΔF/F threshold of 0.15, 14 of the 18 glomeruli imaged from the *Ae. aegypti* AL elicited strong responses to a panel of odorants identified in the *P. obtusata* headspace, DEET (another bioactive odorant), mixtures including the orchid scents and human scent, and the mineral oil (no odor) control. Odorants of the different chemical classes elicited distinct responses in glomeruli (Kruskal-Wallis test: p < 0.001) and glomerular clusters were significantly different in their response (p < 0.0001). (**A**) (Left) 3D reconstruction of the *Ae. aegypti* AL and location of the imaged glomeruli. Glomeruli were assigned names based on (*14*). (Right) Responses of the AM2, AM3, AM4 and AM5 glomeruli to odor stimuli. Bars are the mean ± SEM (n = 7-14 mosquitoes). (**B**) As in A, except for the AL1, AL2 and AC1 glomeruli. (**C**) As in A, except for the AL3, V3, LC1 and LC2 glomeruli. (**D**) As in A, except for the V1, V2, PM1 and PM2 glomeruli.

**Fig. S7.**
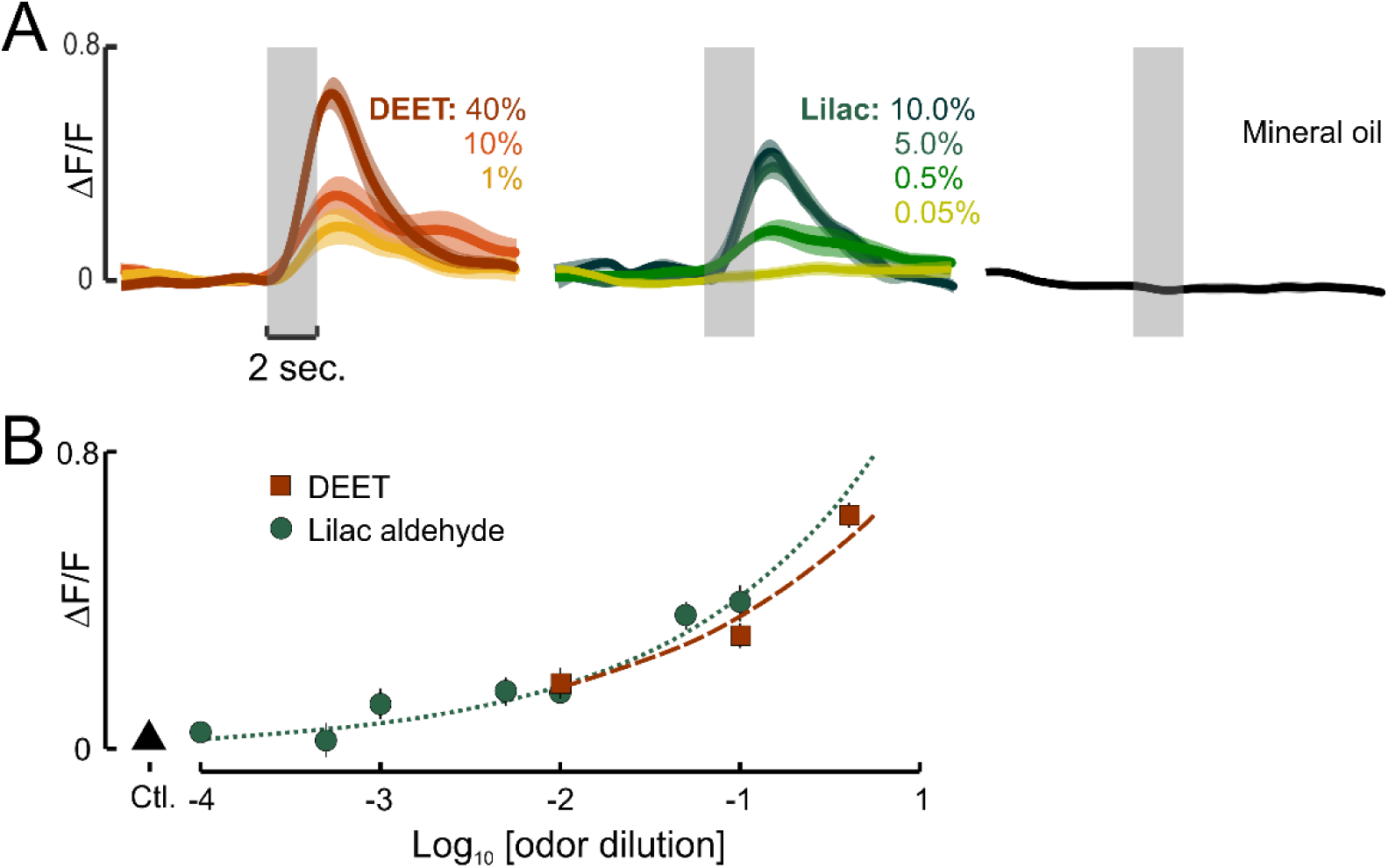
*Ae. aegypti* AM2 responses to lilac aldehyde and DEET at different concentrations. (**A**) ΔF/F time traces for the AM2 glomerulus stimulated at different concentrations of DEET (left, brown), lilac aldehyde (middle, green), and the mineral oil control. Lines are the mean; shaded areas are the SEM (n=4-10 mosquitoes). (**B**) Dose-response curves for AM2 responses to DEET and lilac aldehyde. Both odorants elicited significant increases in response with increasing dose (R^2^≥0.75; p<0.05) and were not significantly different in their model fits (p=0.06)(lilac aldehyde: y = 1.01x^0.39^; DEET: y = 0.77x^0.33^).

**Fig. S8.**
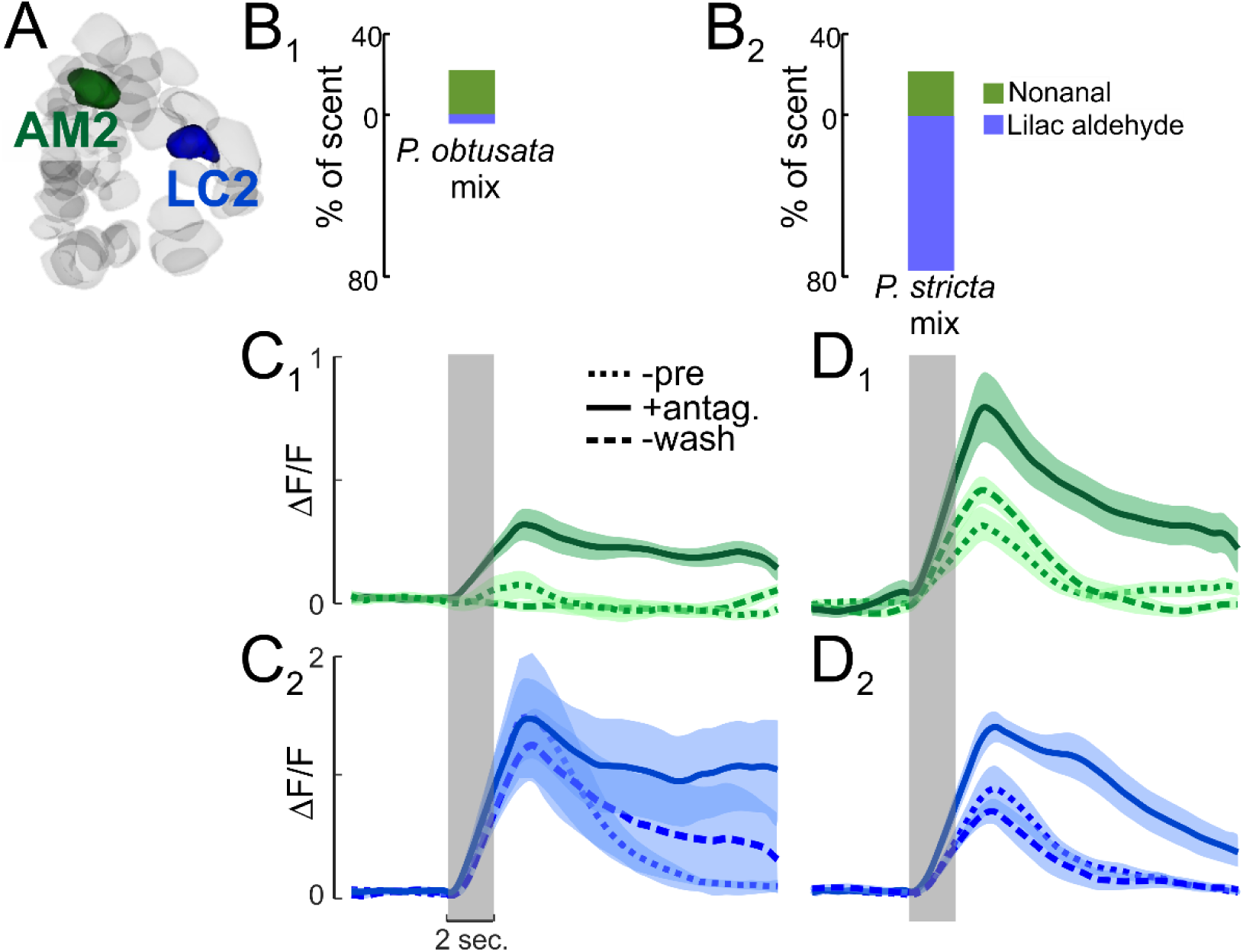
ΔF/F time traces of AM2 and LC2 responses to *P. obtusata* and *P. stricta* mixtures during GABA receptor antagonist application. (**A**) *Ae. aegypti* AL reconstruction showing LC2 (blue) and AM2 (green) glomeruli. (**B**) Ratio of nonanal and lilac aldehyde in the *P. obtusata* (B_1_) and *P. stricta* (B_2_) mixtures. (**C**) AM2 (C_1_, green) and LC2 (C_2_, blue) responses to the *P. obtusata* mixture during (pre)saline superfusion (dotted lines), GABA receptor antagonist application (solid lines), and saline wash (dashed lines). AM2 responses were significantly modified by application of the GABA receptor antagonists (Kruskal-Wallis test: p<0.01), but LC2 responses were not significantly different between pharmacological treatments (Kruskal-Wallis test: p=0.98). Each trace is the mean; areas around the traces are the ± SEM (n=8 stimulations from 4 mosquitoes). (**D**) As in C, but AM2 (D_1_) and LC2 (D_2_) responses to the *P. stricta* mixture during (pre)saline superfusion (dotted lines), GABA receptor antagonist application (solid lines), and saline wash (dashed lines). Both AM2 and LC2 responses were significantly modified by GABA receptor antagonist application (Kruskal-Wallis test: p<0.05).

**Fig. S9.**
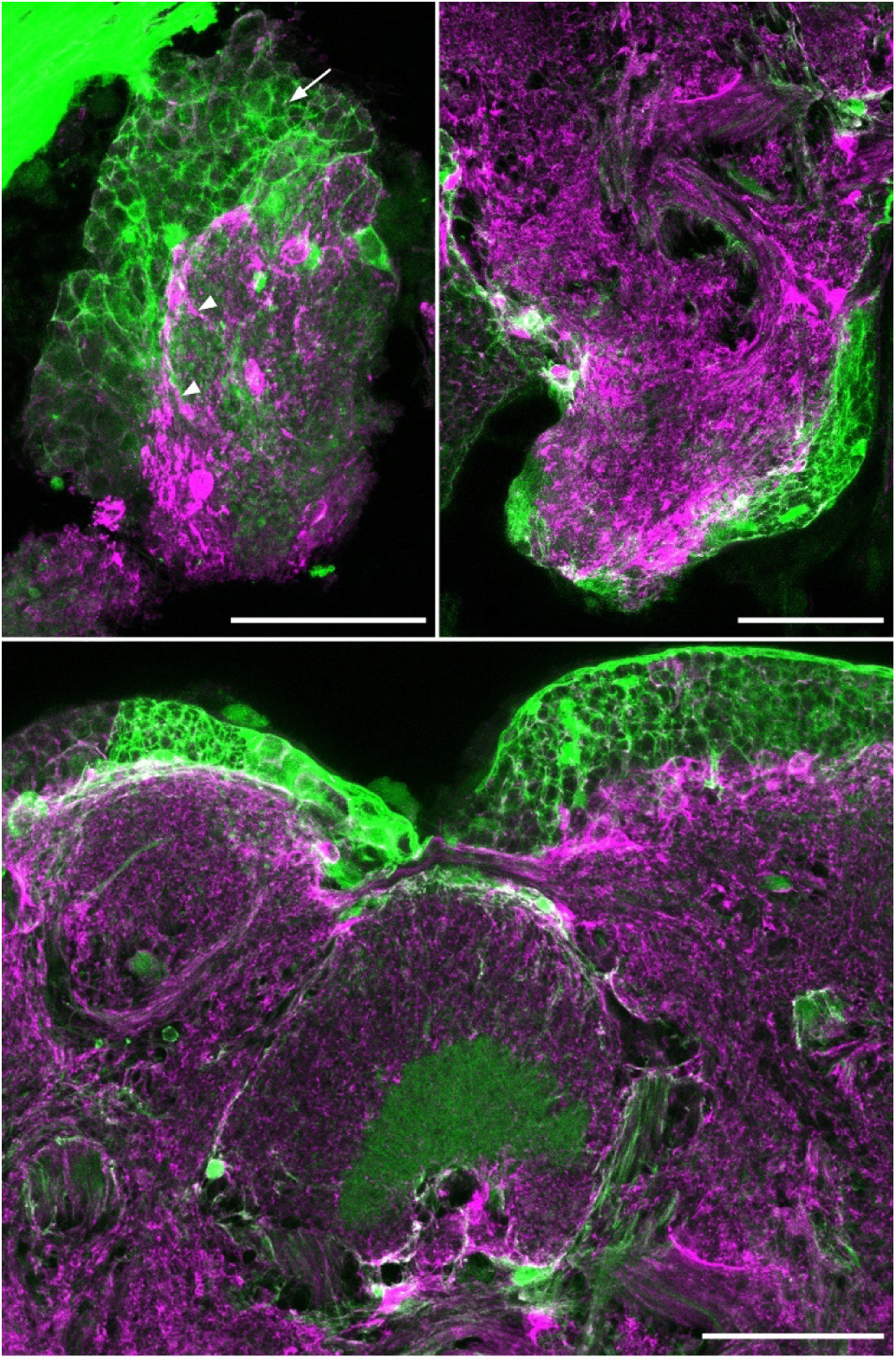
Confocal images brain sections stained for GFP (GCaMP) and glutamate synthase (glia). Confocal images of brain sections from *PUb-GCaMP6s Ae. aegypti*. (*Upper left*) In the AL, GFP immunofluorescence (green) reveals expression of GCaMP6s, which does not overlap with glia, labeled with antisera against glutamate synthase (GS, magenta). Arrow in upper left panel indicates neuronal cell bodies in the lateral antennal lobe cluster. (*upper right*) The Mushroom Body Calyx; and (*Lower center*) the Central Complex. Scale bars are 100 µm.

**Table S1.**
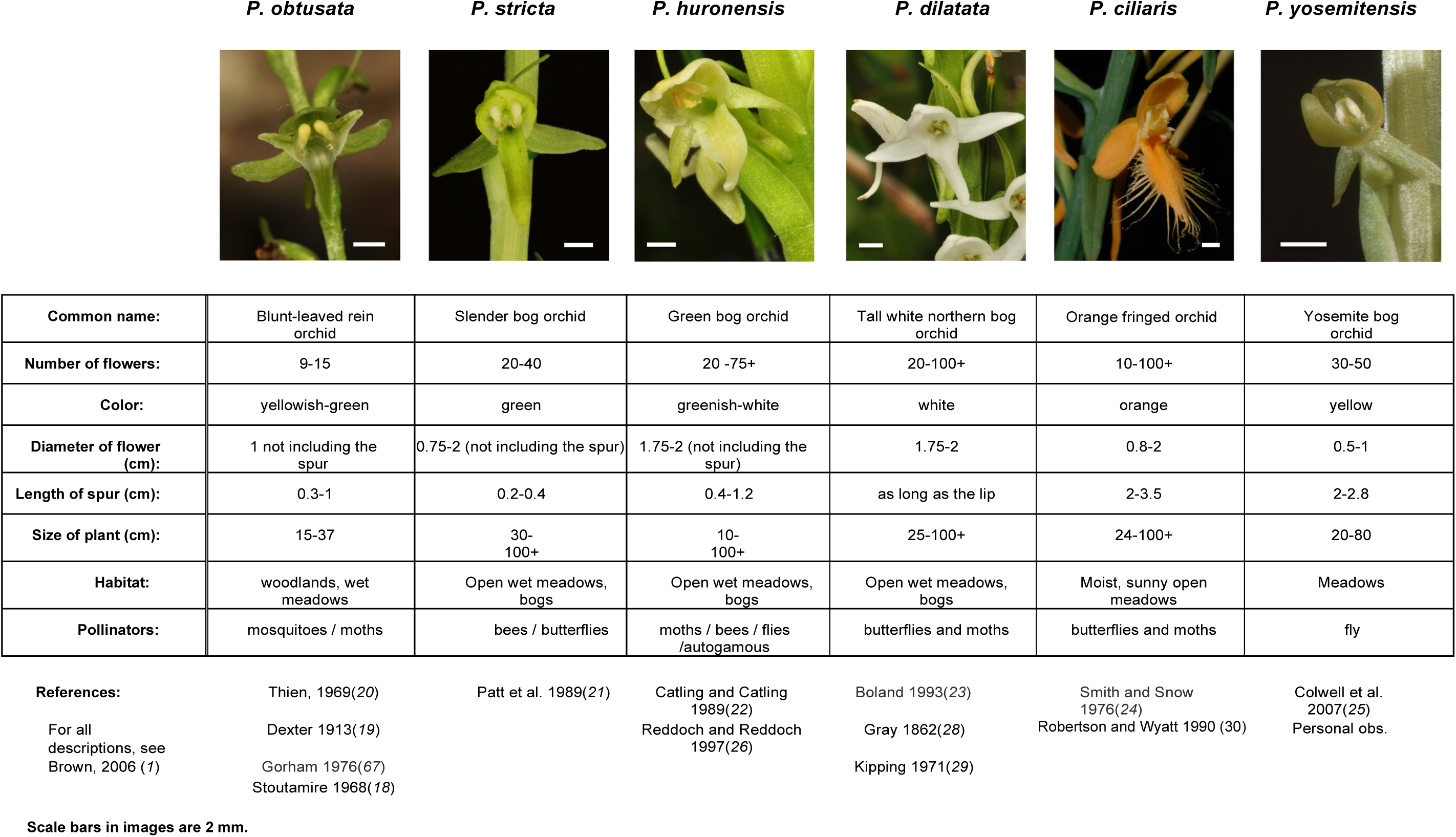
*Platanthera* spp. morphological traits and pollinators.

**Table S2.** Mosquito species captured in the field and numbers of individuals found with attached pollinia.

| Mosquito genus | Mosquito species | Individuals caught with pollinia attached |
| --- | --- | --- |
| ***Aedes*** | *Ae. cinereus* | 1 |
| ***Aedes* (Subgenus *Ochlerotatus*)** | *Ae. communis* | 24 |
|  | *Ae. increpitus* | 9 |
|  | *Ae. fitchii* | - |
|  | *Ae. dianteus* | - |
|  | *Ae. canadensis canadensis* | 1 |
|  | *Ae. implicatus* | 1 |
|  | *Ae. sp.* | 3 |
| ***Culiseta*** | *Cs. impatiens* | - |
|  | *Cs. incidens* | - |
| ***Anopheles*** | *An. freeborni* | - |
| ***Culex*** | *Cx. pipiens* | - |

**Table S3.**
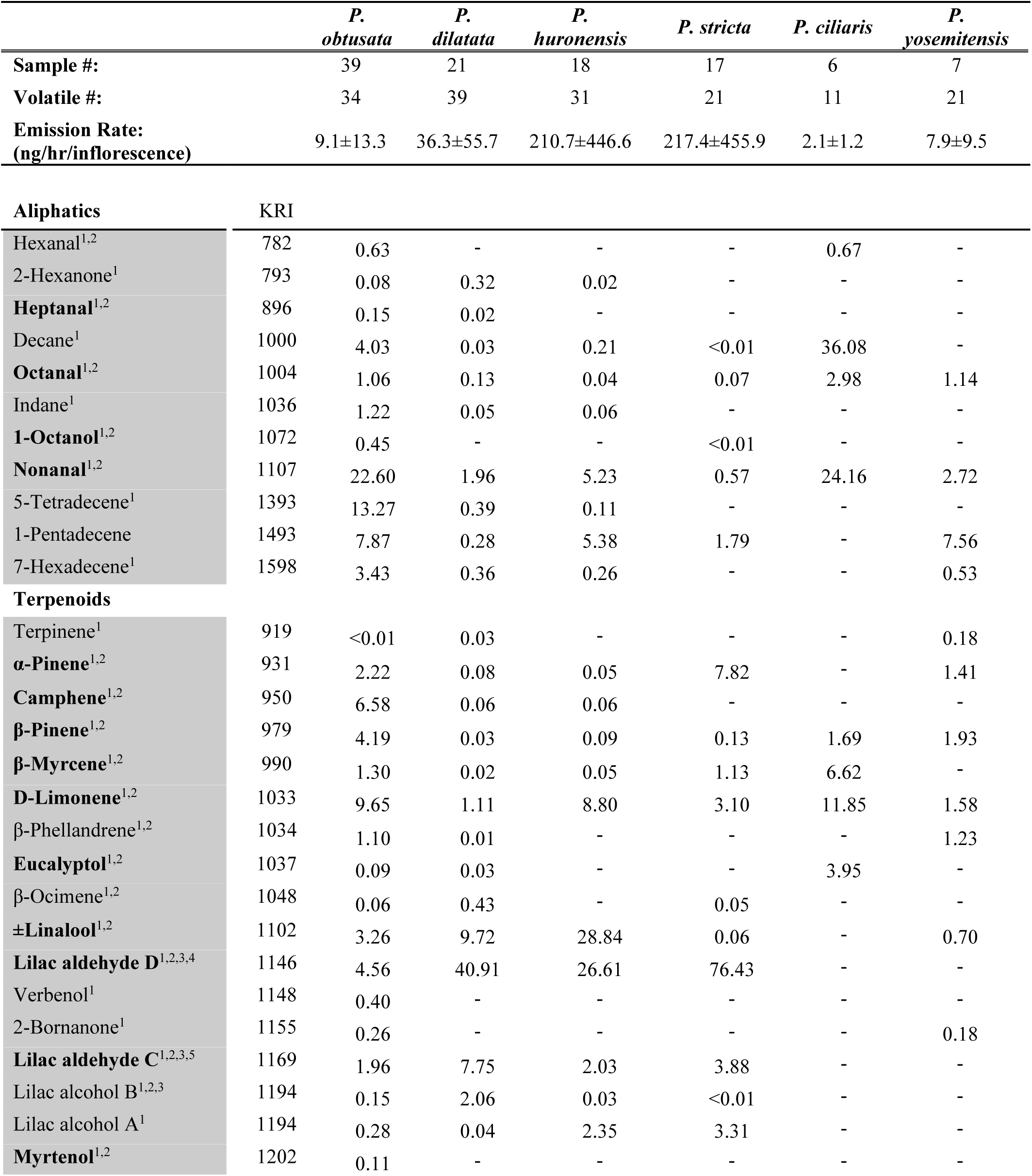
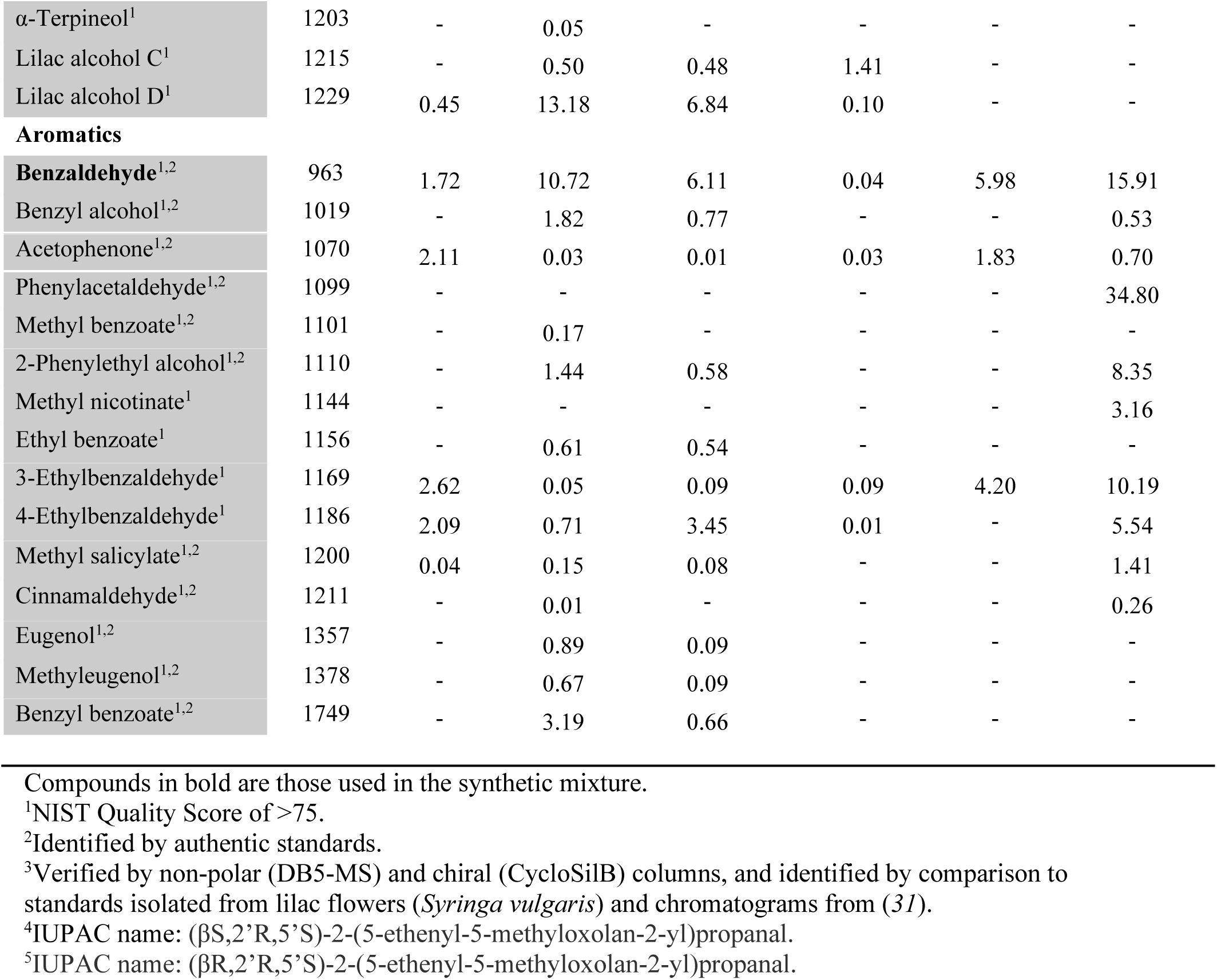
Composition and emission rates of the *Platanthera* orchid scents. The values for the volatile compounds in the scent of each orchid species are presented as percentages. Emission rates are the mean ± SD.

